# Contralesional activity reflects compensation, while brainstem detour pathways support skilled motor recovery after stroke

**DOI:** 10.1101/2025.11.24.690099

**Authors:** Matteo Panzeri, Nina Thiem, Christian Hoffmann, Ulrike Schillinger, Nirav Barapatre, Simon Musall, Fritjof Helmchen, Anna-Sophia Wahl

**Affiliations:** Brain Research Institute, University of Zurich, Winterthurerstrasse 190, 8057 Zurich, Switzerland; Neuroscience Center Zurich (ZNZ), University of Zurich, Zurich, Switzerland; Institute for Stroke and Dementia Research (ISD), LMU University Hospital, LMU Munich, Feodor-Lynen-Strasse 17, 81377 Munich, Germany; Department of Neuroanatomy, Institute of Anatomy, Ludwigs-Maximilians-University, Pettikoferstrasse 11, 80336 Munich, Germany; Institute of Biological Information Processing, Department for Bioelectronics (IBI-3), Forschungszentrum Jülich, Jülich, Germany; Department of Systems Neurophysiology, Institute for Zoology, RWTH Aachen University, Aachen, Germany; Faculty of Medicine, Institute of Experimental Epileptology and Cognition Research, University of Bonn, Bonn, Germany; University Research Priority Program (URPP), Adaptive Brain Circuits in Development and Learning, University of Zurich, Zurich, Switzerland

**Keywords:** large strokes, cortical and subcortical reorganization, brainstem nuclei, motor centers brainstem, modeling of brain-wide reorganization

## Abstract

Large strokes frequently result in lasting motor deficits and trigger extensive reorganization within the brain and spinal cord. Altered neuronal activity in the contralesional hemisphere has been documented in both humans and rodent models, yet its role in functional recovery versus maladaptation remains unresolved. Here, we used chronic wide-field calcium imaging to monitor bilateral cortical activity in mice performing a skilled reach-to-grasp task before and days to weeks after stroke. Strokes produced persistent fine motor impairments, which were only partly alleviated by intensive rehabilitative training. While cortical activity was in particular suppressed in the ipsilesional cortex after stroke, training promoted sustained increases in contralesional sensorimotor activity. However, ridge regression analysis of neural and behavioral data indicated that this activity largely reflected compensatory use of the intact paw rather than recovery of the impaired forelimb. Axonal tracing nevertheless revealed enhanced projections from contralesional motor cortex to ipsilesional brainstem nuclei - including the midbrain and pontine reticular nucleus - specifically in trained animals. These findings identify a rehabilitation-induced corticofugal pathway supporting motor recovery, highlighting a target for neuromodulation strategies in chronic post-stroke impairment.

## Introduction

The contribution of the contralesional, intact hemisphere to post-stroke recovery has remained a subject of debate across decades of basic and clinical research. It is now almost canonical knowledge that, following small cortical strokes, functional remapping primarily occurs within the peri-infarct cortex, whereas larger lesions elicit more extensive structural and functional reorganization in the contralesional hemisphere^1^. Nevertheless, the extent to which circuit-level rewiring within the contralesional hemisphere supports the restoration of lost or impaired functions - particularly sensorimotor capabilities - remains incompletely understood.

Chollet *et al.*^2^ first reported increased cerebral blood flow in the contralesional hemisphere of stroke patients who had recovered from hemiplegic hand movements. Subsequent PET^3^ and fMRI studies^4–7^, as well as meta-analyses^8^, consistently confirmed that enhanced activity within the contralesional primary motor cortex (M1), bilateral ventral premotor cortex, and supplementary motor area (SMA) is detected in strokes affecting the motor system. However, such contralesional activation has also been linked to poorer motor outcomes^9,10^, suggesting that successful recovery may depend on the normalization of pathologically elevated interhemispheric activity over time^6,8,10,11^.

An alternative view posits that contralesional hyperactivity may actively hinder recovery, as excessive neuronal activity in the intact hemisphere could impose maladaptive interhemispheric inhibition on ipsilesional motor areas, thereby constraining the reorganization of residual motor circuits^4,12^. Attempts to reconcile these seemingly contradictory findings, where both stimulation^13^ and inhibition of the contralesional hemisphere^14–16^ have been shown to improve motor outcomes, have led to the proposal that contralesional hyperactivity represents a dynamic, time-dependent process. In this framework, enhanced activity in the contralesional hemisphere may initially facilitate recovery through cross-cortical and subcortical reorganization, but its persistence beyond the early post-stroke phase may become maladaptive, requiring subsequent downregulation for optimal functional restoration.

Here we introduce an experimental paradigm that enables the longitudinal assessment of fine motor function before and after stroke in mice, combining behavioral analysis with mesoscale mapping of cortical reorganization through chronic wide-field calcium imaging. Using a skilled forelimb grasping task, we observed increased neuronal activity within the contralesional, intact hemisphere, particularly in regions involved in motor planning and execution. This enhancement was most pronounced in animals that underwent intensive motor training following stroke. However, quantitative modeling of movement-related neural activity, incorporating uninstructed body movements of the unaffected forelimb, as well as task-related variables, revealed that contralesional hyperactivity was not directly associated with improved motor performance of the impaired paw. Instead, it primarily reflected compensatory adjustments of the intact limb during skilled grasping. In contrast, anatomical tracing from the contralesional cortex uncovered extensive corticofugal sprouting forming alternative, “detour” pathways through ipsilesional brainstem nuclei. These newly established projections were strongly associated with recovery in skilled grasping performance after large cortical strokes, suggesting that structural reorganization, rather than contralesional hyperactivity per se, underlies effective functional restoration.

## Results

### Large strokes impair skilled motor function, which can be in part reversed by intensive physical training

To study cortical reorganization in relation to fine motor behavior, we developed a lever-pressing task for head-fixed skilled motor assessment combined with chronic widefield calcium imaging of cortical activity in the healthy condition and several weeks after stroke (Figure 1A). Transgenic mice expressing the calcium indicator GCaMP6f (GP5.17 Thy1-GCaMP6f) were trained to reach and pull a joystick after an auditory cue (8 kHz, Supplementary Video 1). Upon successfully producing a sufficiently large joystick deflection (>10 mm) within a 5-s response window, animals received a sweetened water reward. We used high-speed camera recordings (150 Hz, Figure 1A, B) for detailed kinematic analysis of grasping movements of the task limb and the supporting limb while simultaneously performing widefield calcium imaging of neuronal population activity in both hemispheres through the intact skull. After an initial handling and training phase of 3 weeks (Figure 1C) animals reached expert performance in the task (> 80% of rewarded trials for 3 consecutive sessions). We then randomized animals in Sham (n = 5 mice), Stroke (n = 9 mice), and Rehab (n = 7 mice) groups (Figure 1C).

**Figure 1.**
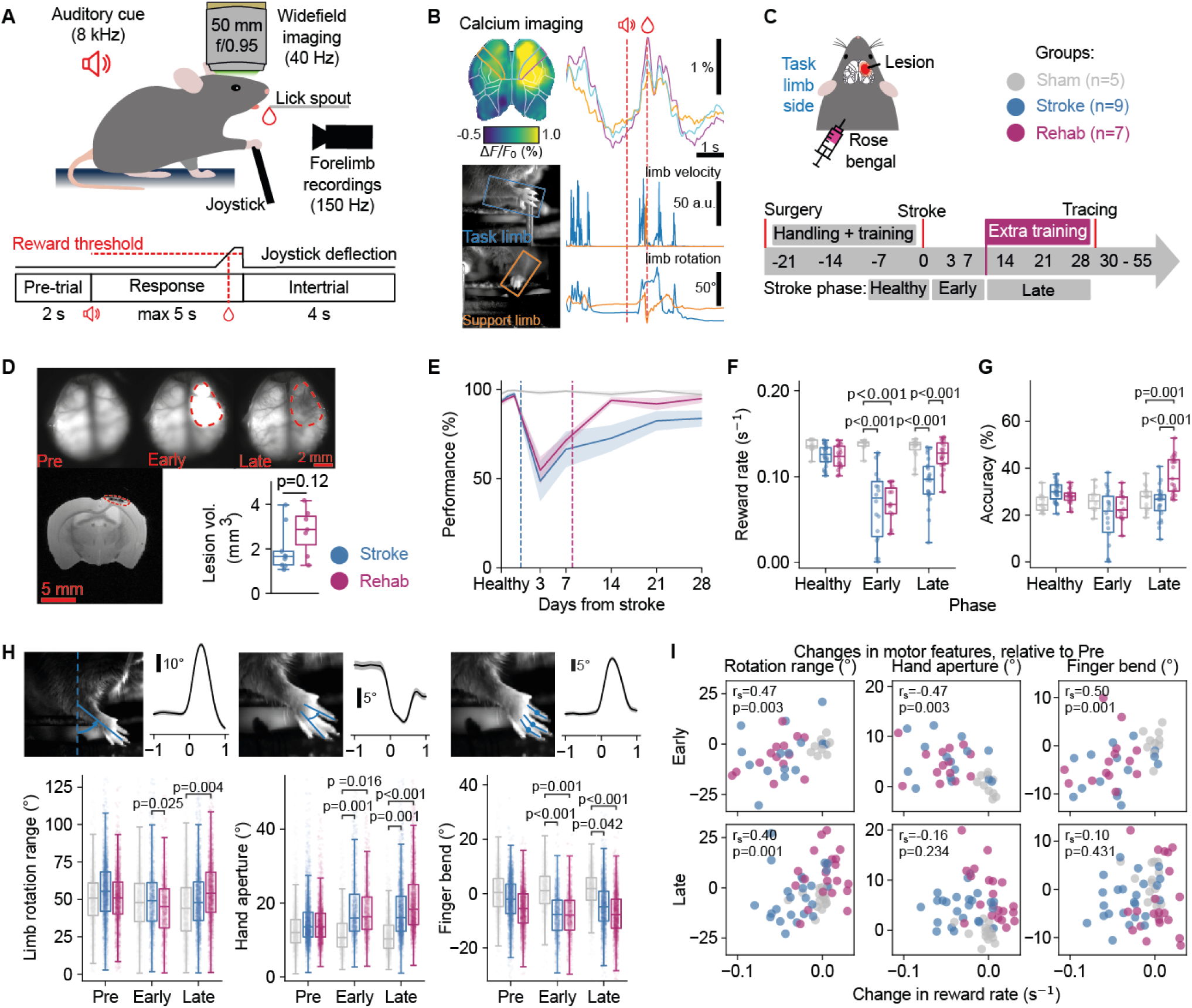
Large strokes impair skilled motor function, which can be in part reversed by intensive physical training. **A.** Experimental setup: mice were trained to pull a joystick and receive a water reward following an auditory cue. During the task, neuronal activity and limb movements were recorded. **B.** Representative data from an experimental session. Top: widefield spatial activity map, and transients from highlighted ROIs. Bottom: example camera views of task and support limbs, and velocity and limb rotation traces for each limb. **C.** Top: photothrombotic stroke schematic and experimental cohorts. Bottom: experimental timeline. **D.** Top: example widefield imaging field of view at different experimental phases. Red dotted line highlights the ischemic lesion. Bottom left: T2-weighted MRI scan used for quantifying lesion sizes. Bottom right: lesion size comparison across cohorts (n= 9 Stroke and n= 7 Rehab mice). **E.** Task performance across time, averaged across mice by experimental cohorts. Shaded area represents s.e.m.. **F.** Comparison between cohorts of the reward rate, defined as the task performance normalized by session length, at different stroke timepoints. Datapoints represent individual sessions (n=163 sessions, n=21 mice). **G.** Comparison between cohorts of task accuracy, defined as the percentage of rewarded grasps to total grasps. Datapoints represent individual sessions (n=163 sessions, n=21 mice). **H.** Comparison of experimental cohorts’ different fine motor features during rewarded grasps. For all quantifications, each datapoint represents one single grasp (n=14115 grasps, n=21 mice). Left: (top) rotation range (max-min rotation over a grasp) example of time course aligned to movement onset, and quantifications (bottom). Middle: hand aperture, quantification represents the mean over time from start to end of each grasp. Right: finger bending, quantification represents the mean over time from start to end of the grasp. **I.** Spearman correlations between session-averaged fine motor features and reward rate. All datapoints represent individual sessions (n=99 sessions, n=21 mice), and all data is normalized per-mouse by subtracting the pre-stroke baseline median. In **D.**, statistical comparison was carried out using a two-tailed independent samples t-test. In **E**-**H,** statistical comparisons were evaluated using linear mixed-effect models, and contrasts were evaluated on the estimated marginal means between groups. Reported p-values are adjusted for multiple comparisons (post-hoc Bonferroni correction for each model).

After introducing a large photothrombotic stroke (see methods)^13,17^, destroying the sensorimotor cortex corresponding to the task limb trained in the lever-pressing task, all animals were re-assessed for their grasping performance at day 3 and 7 after stroke (early phase). We continued (day 8-28, late phase) combining weekly assessment in the grasping task during widefield calcium imaging for up to 4 weeks after stroke (Figure 1C). We trained the animals in the Rehab group as a form of motor rehabilitation training in the lever-pressing task for 4 additional training sessions per week (100 reaching attempts each), starting on day 8 after stroke (Figure 1C). Ex-vivo MRI did not show a significant difference of stroke lesion size between animals with and without rehabilitative training (lesion volume, Stroke: 1.96 ± 0.31 mm^3^, Rehab: 2.80 ± 0.36 mm^3^, p = 0.124, Figure 1D).

We first examined how the stroke affected the performance in the lever-pressing task (Figure 1E). UMAP embedding of all movements before stroke had already revealed distinct kinematic clusters of successful “rewarded” grasps, “missed grasps” and “other movements” (Supplementary Figure 1A) with noticeable differences in fine motor features of the task paw such as limb rotation, hand aperture and finger bend (Supplementary Figure 1B, see Methods for definitions). To detect motor recovery after stroke, we solely focused on rewarded grasps with successful deflection of the joystick above the reward threshold. We found a significant and comparable drop in the rate of rewarded grasps for both stroke groups relative to Sham in the early-post stroke phase (Stroke -Sham: −0.070s^−1^, 95% CI [−0.094,−0.047], p<0.001; Rehab - Sham: −0.068s^−1^, 95% CI [−0.093,−0.044], p<0.001; Rehab – Stroke: 0.002s^−1^, 95% CI [−0.019,0.023], p=1; linear mixed model, Figure 1F). In the late post-stroke phase only the Rehab group, not the Stroke group, recovered to levels comparable to the Sham group (Stroke - Sham:-0.038s^−1^, 95%CI [−0.060,−0.017], p<0.001; Rehab - Sham: −0.007s^−1^, 95%CI [−0.030,0.015],p=1; Rehab - Stroke: 0.031s^−1^, 95% CI [0.012, 0.017], p<0.001; linear mixed model, Figure 1F). Furthermore, mice that underwent physical training displayed a higher grasping accuracy (see Methods) in the late post-stroke phase, relative to both Sham and Stroke groups (Rehab - Sham: 9.85%, 95%CI [3.31,16.4], p=0.001; Rehab - Stroke: 11.5%, 95%CI [5.83,17.3], p<0.001; linear mixed model, Figure 1G).

A detailed analysis of grasping behavior using automatized kinematic analysis with DeepLabCut^18^, detected a positive effect of the training in the Rehab group only for proximally driven limb rotations (Rehab - Sham: 11.9°, 95%CI [2.92,21.0], p=0.004; linear mixed model, Figure 1H left). At the same time, a chronic motor deficit persisted for distal motor parameters not only in the Stroke, but also for the Rehab group. We detected an impaired aperture of the hand (Late Stroke - Sham: 7.27°, 95%CI [2.32,12.2], p=0.001; Late Rehab - Sham: 9.20°, 95%CI [4.03,14.4],p<0.01; linear mixed model, Figure 1H middle) and a decreased capacity to bend fingers (Late Stroke-Sham: −5.97°, 95%CI [−11.8,−0.15], p=0.042; Late Rehab -Stroke: −9.26°, 95%CI [−15.3,−3.19], p<0.001; linear mixed model, Figure 1H right). Changes in fine motor parameters correlated with the rate of rewarded trials in the early phase (rotation range: Spearman r=0.47, p=0.003; hand aperture: Spearman r=−0.47, p=.0003; finger bend: Spearman r: 0.50, p=.001, linear mixed model, Figure 1I). But only the limb rotation was correlated to improved performance in the lever-pressing task, indicated as increase in rewarded trials (rotation range: Spearman r:0.40, p=0.001; linear mixed model, Figure 1I), suggesting that in the late phase, mice were able to perform the task despite chronic distal motor impairment. Inspection of the same fine motor metrics for support limb movements, as well as quantifications of movement rates, did not show major modifications in the use of the support limb (Sham versus Stroke versus Rehab group, Supplementary Figure 1C, D) over the course of the experiment. There was also no relation between stroke lesion size and distinct motor kinematics early or late after stroke (Supplementary Figure 1E, F).

### Contralesional cortical activity is enhanced in mice with intensive rehabilitative training

We next asked how the pattern of cortical reorganization after stroke related to the distinct motor outcome in the different treatment groups examined. We performed chronic widefield calcium imaging in the ipsilesional and contralesional cortices, related to task limb and support limb, respectively, before and after stroke (Figure 2A). To analyze task-related cortical activity patterns over time, we computed average calcium signals aligned to the onset of rewarded grasps. We first aligned the data to the Allen Brain Atlas^19^, and computed pre-stroke cortical activity maps, as a baseline to which cortical activity changes were compared (Figure 2B - average of all pre-stroke sessions). As variables of interest, we computed the average calcium signals in regions of interest (ROIs) representing Allen Brain Atlas regions. We compared the mean activity over a response window of 1 s after the onset of rewarded grasps to the baseline (Figure 2C - shaded area). Compared to the Sham group, we found decreased post-stroke cortical responses during rewarded grasps in the ipsilesional primary motor cortex (M1) in both stroke groups in the early (Stroke-Sham: −0.611%, 95%CI [−0.885,−0.336], p<0.001; Rehab - Sham: −0.732%, 95%CI [−1.02,−0.445], p<0.001; linear mixed model, Figure 2D top) and late post-stroke phases (Stroke-Sham: −0.484%, 95%CI [−0.752,−0.216], p=0.004; Rehab - Stroke: −0.441%, 95%CI [−0.722,−0.161], p=0.013; linear mixed model, Figure 2D top). In contrast, we measured increased activity in the late post-stroke phase in contralesional M1, which was only significant in mice with intense rehabilitative training (Rehab versus Stroke group: 0.360%, 95% CI [0.195, 0.523], p<0.001; linear mixed model, Figure 2D). We repeated the same analysis over all the ROIs in the field of view, subtracting cortical responses between groups and found enhanced cortical activity in the rehabilitation group for all sensorimotor and premotor areas compared to Sham or Stroke animals without further motor training, particularly in the chronic, late stage after stroke (Figure 2E). The enhanced cortical activity in the contralesional hemisphere was positively correlated to the rate of successfully performed lever-pressing grasps (early: Spearman r=0.341, p=0.029; late: Spearman r=0.374, p=0.003; Figure 2F left), but negatively correlated with fine motor outcome parameters such as finger bending in the chronic stage after stroke (late, Spearman r=−0.418, p=0.001; Figure 2F right). Cortical activity in ipsilesional M1 was only transiently correlated with an improved success rate of rewarded grasps, while there was no relation to fine motor features (Supplementary Figure 2A-B, Spearman correlation). There was also no clear influence of stroke lesion volume on either ipsilateral or contralateral M1 activity early or late after stroke (Supplementary Figure 2C). Furthermore, when limiting the set of rewarded grasps by only including grasps with similar features compared to their pre-stroke template (Supplementary Figure 3A), we still observed an increase in contralesional activity for the Rehab group, suggesting that the contralesional activity increase could not be explained by changes in the trajectory of the task limb (Supplementary Figure 3B). Likewise, restricting the set of rewarded grasps including only grasps accompanied by similar support-limb movements also retained the increase in contralesional activity for the Rehab group (Supplementary Figure 3C-D).

**Figure 2.**
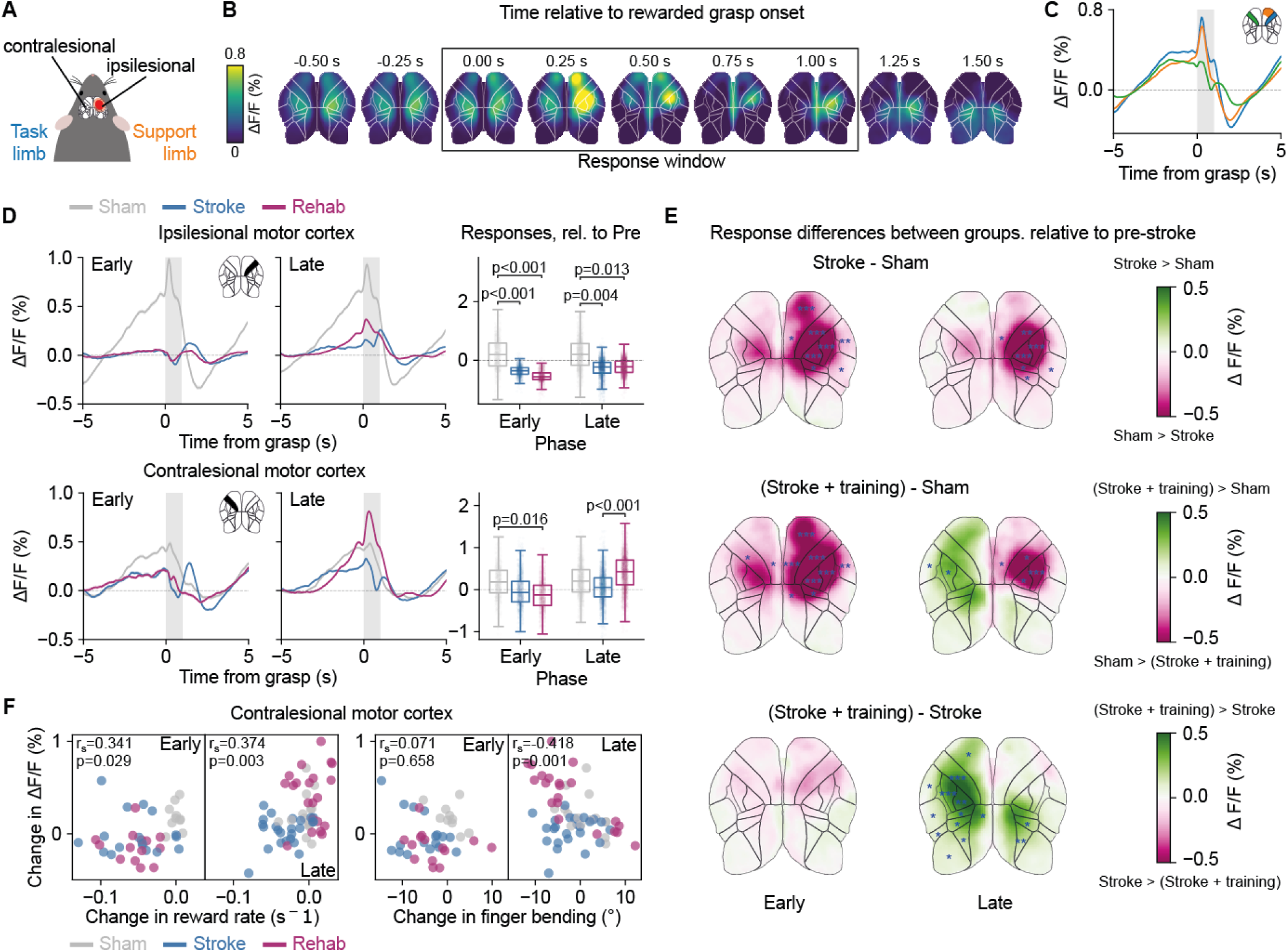
Contralesional cortical activity is enhanced in mice with intensive rehabilitative training. **A.** Schematic defining the ipsilesional and contralesional cortex locations relative to the task and support limbs. In all plots, the ipsilesional hemisphere is shown on the right and the contralesional hemisphere on the left. **B.** Average calcium activity timecourses in the pre-stroke condition, aligned to the onset of rewarded grasps. Averaged across all pre-stroke sessions (n=5772 grasps, n=61 sessions, n=21 mice). Black box denotes the response window, across which activity is averaged, for the purpose of statistical comparisons. **C.** Average timecourses over all pre-stroke rewarded grasps (mean and s.e.m.), for highlighted areas. Areas include ipsilesional and contralesional M1 (blue and green) and the anterior half of ipsilesional M2 (orange). Shaded bar represents the response window. (n=5772 grasps, N=21 mice). **D.** Top: (left) average timecourses of the ipsilesional M1, by experimental cohort, in the early and late post-stroke phases. (right) quantification of changes relative to pre-stroke baseline. Datapoints represent mean over the response window for individual grasps (n=8112 grasps, n=21 mice). **E.** Differences in average response-window between experimental groups at different timepoints. Top: Stroke – Sham, middle: (Stroke + training) – Sham, bottom: (Stroke + training) – Stroke. Overlaid in blue are the results of statistical comparisons between ROI-averaged responses (Sham: n=2417 grasps; Stroke: n=2893 grasps; Rehab: n=2802 grasps). **F.** Spearman correlations between changes for the contralesional M1, and the reward rate (left), or finger bending (right) for all groups. Datapoints represent individual sessions (averaged over grasps); values are relative to pre-stroke baseline (n=101 sessions, n=21 mice). In **D.** and **E.** statistical comparisons were evaluated by fitting linear mixed models (one model per ROI) and p-values were adjusted for multiple comparisons across groups, time and rois by controlling the false discovery rate (Benjamini-Hochberg correction). In **E.,** blue asterisks indicate significances for the corresponding ROI: *p<0.05, **p<0.01, *** p<0.001.

### Variance of neuronal activity explained by behavioral components using ridge regression models

Given that contralesional cortical activity was particularly elevated in animals undergoing intensive motor training after stroke, we sought to determine whether this enhanced activity was directly related to motor recovery, or instead reflected uninstructed movements and compensatory strategies supporting task performance. To disentangle these possibilities, we implemented a ridge regression model designed to quantify the contribution of distinct behavioral components - including movements of the task and support limbs, licking during reward consumption, and sensory responses to the auditory cue - to the mesoscale cortical activity measured via wide-field calcium imaging (Figure 3A). We linearly partitioned the ΔF/F predicted by the full model into the contribution of each behavioral variable. In the pre-stroke condition, the model reliably predicted cortical activity aligned to rewarded grasp onsets in both hemispheres. In particular, task-limb movements were the dominant contributors of activity within the corresponding ipsilesional M1 region, whereas both task- and support-limb movements contributed substantially to contralesional M1 activity (Figure 3B). Extending this approach, we reconstructed spatial activity time courses across all rewarded pre-stroke grasping trials, allowing the decomposition of overall cortical activity into distinct behavioral components (Figure 3C). Spatial maps of uniquely explained variance (Figure 3D) further demonstrated that individual behavioral variables such as the support and task limb could be very well located in the respective motor cortex areas. Furthermore, spatial maps of uniquely explained variances for other behavioral components such as auditory cue processing, selectively accounted for activity within functionally relevant cortical regions, including the auditory cortex. Additionally, we show that our models correctly replicate the post-stroke increase of contralesional activity, by plotting cortical response maps akin to figure 2E, but using the full model predictions (Supplementary Figure 4A). Together, these analyses validate our model’s capacity to resolve the behavioral determinants underlying cortical population dynamics across distributed motor and sensory areas.

**Figure 3.**
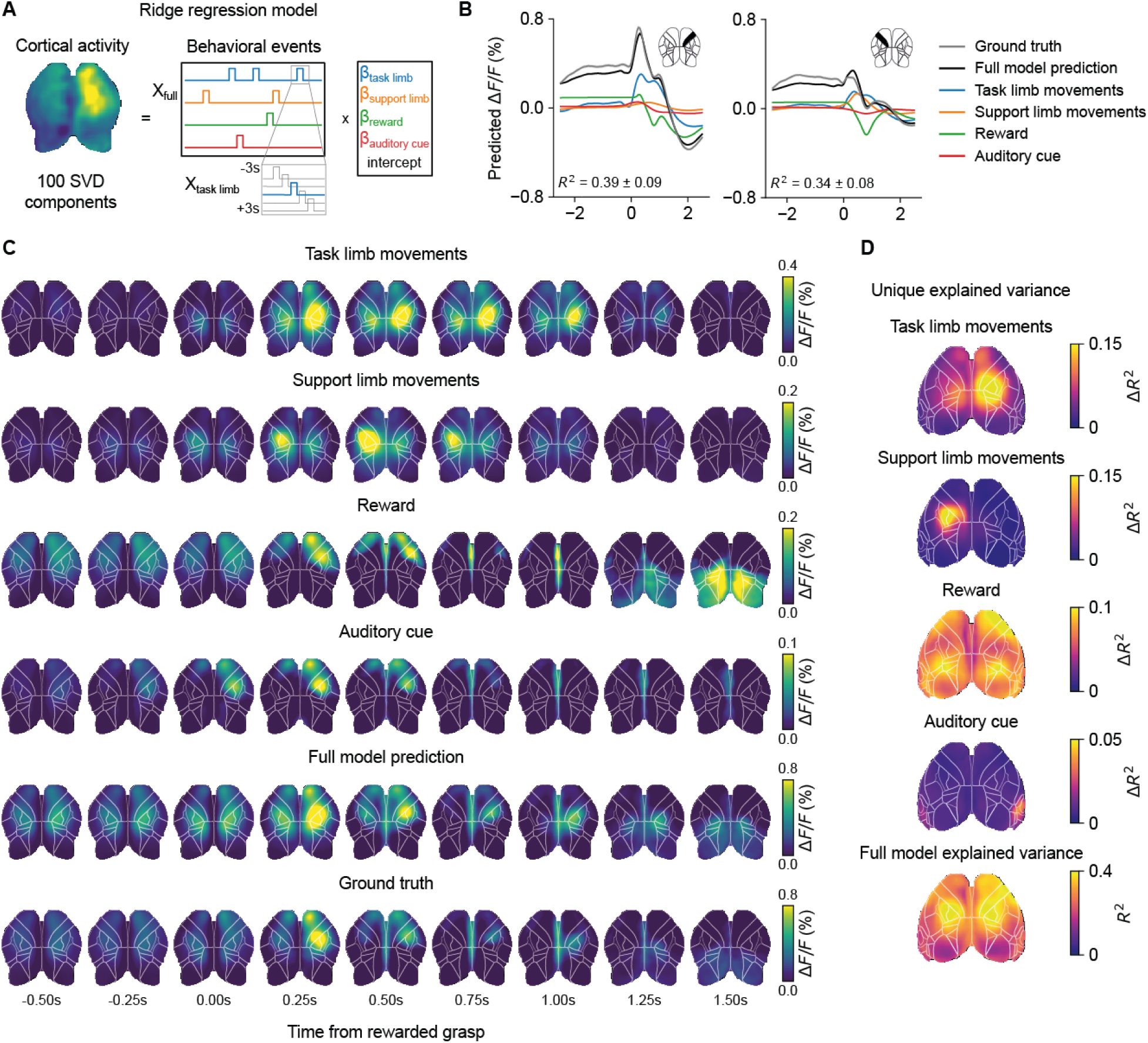
Variance of neuronal activity explained by behavioral components using ridge regression models. **A.** Schematic of the ridge regression model. Neuronal activity (top 100 SVD components) is modeled as a linear combination of binarized behavioral variables, plus time-lagged versions. **B.** Average across all pre-stroke rewarded grasps (n=5772 grasps, n=61 sessions, n=21 mice), partitioned into behavioral contributions. Left: ipsilesional M1, right: contralesional M1. Inset text shows the average explained variance (*R*^2^, +/− standard deviation) of the full model, for the represented ROI. **C.** Average spatial activity timecourses over all pre-stroke rewarded grasps, partitioned into behavioral contributions. **D.** Spatial unique explained variance maps (Δ*R*^2^) for each behavioral variable. Averages across pre-stroke sessions (n=61 sessions). Bottom: spatial *R*^2^ for the full model prediction.

### Increased contralesional activity during rewarded grasps reflects support limb representation rather than reorganization for impaired limb recovery

We next applied our ridge regression model to predict cortical activity driven by distinct behavioral components comparing changes in predicted ΔF/F between experimental conditions (Sham vs. Stroke, Sham vs. Rehab, and Stroke vs. Rehab) at early and late post-stroke stages (Figure 4). Predicted activity in ipsilesional sensorimotor regions corresponding to the task limb was reduced late after stroke (Ipsilesional M1, late; Stroke-Sham: −0.139%, 95%CI [−0.237,−0.041], p=0.049; Rehab -Sham: −0.145%, 95%CI [−0.247,−0.043], p=0.049; linear mixed models, Figure 4A) and not significantly increased in the contralesional sensorimotor regions (Contralesional M1, late; Stroke-Sham:0.001%, 95%CI [−0.070,0.067], p=0.99; Rehab - Stroke: 0.011%, 95%CI [−0.061,0.083], p=0.99; linear mixed models, Figure 4A). By contrast, increased contralesional cortical activity in the late phase - particularly in animals receiving intensive motor training - was predominantly explained by movements of the support limb (Contralesional M1, late; Rehab - Sham: 0.075%, 95%CI [0.035,0.116], p=0.013; linear mixed models, Figure 4B). Predicted activity maps associated with other task variables such as auditory cues or reward processing, revealed an increase, albeit not significant, for mice receiving intensive physical training (Contralesional M1, late; (Rehab - Stroke: 0.196%, 95%CI [0.067, 0.325], p=0.072; linear mixed models, Figure 4C).

**Figure 4.**
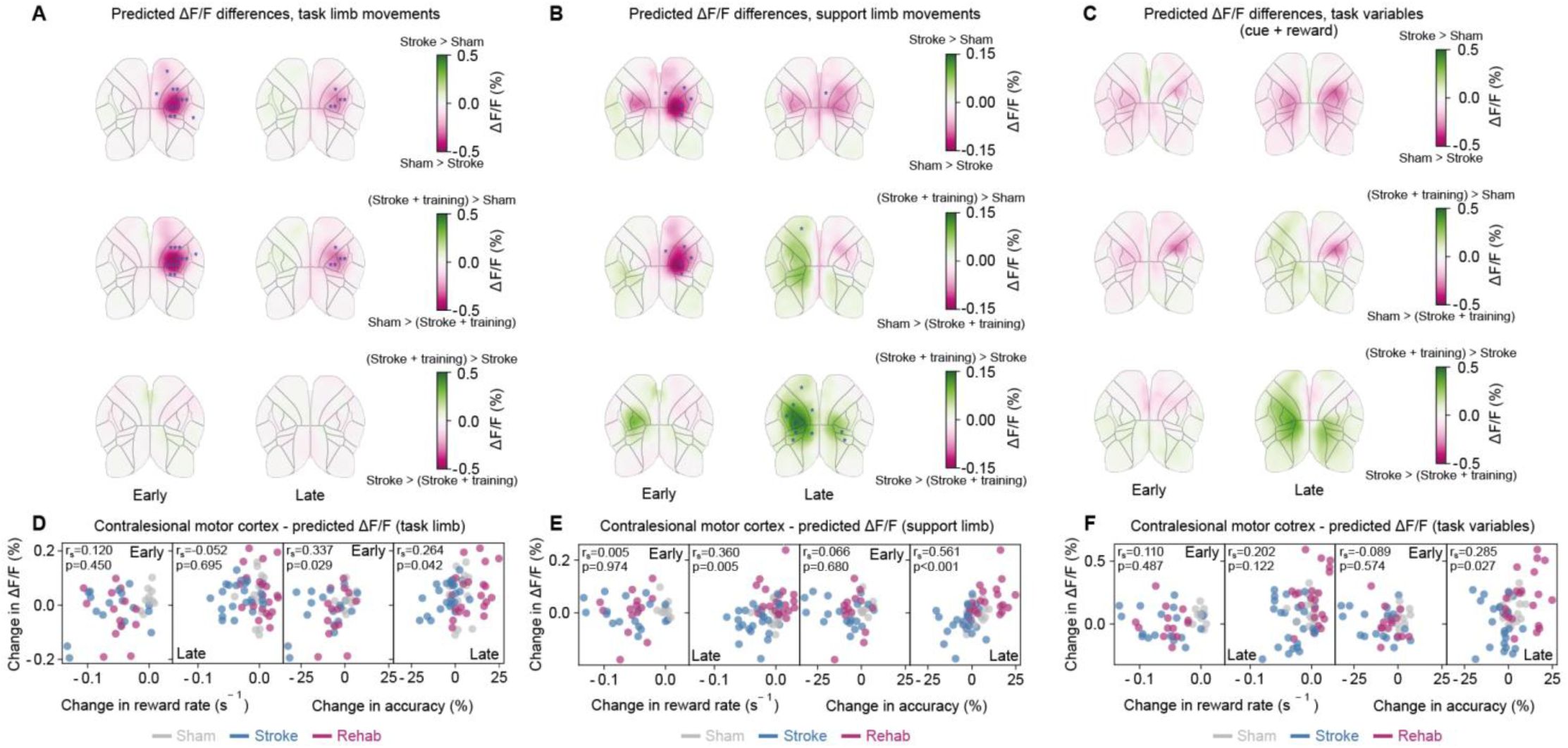
Increased contralesional activity during rewarded grasps reflects support limb representation rather than reorganization for impaired limb recovery. **A.** Differences in predicted by task limb movements, between cohorts at different post-stroke phases. Datapoints are response windows averaged (0.0s −1.0s during a grasp), over individual grasps (n=8247 grasps, n=21 mice). **B.** and **C.** are the same as **A.** but generated using predicted by support limb movements and task variables (auditory cue + reward), respectively. **D.-F.** Spearman correlation between contralesional cortex (task-limb driven in **D.** or support limb driven in **E.** or task variables driven in **F.**) and reward rate and accuracy. Datapoints represent session-averaged (n=102 sessions, n=21 mice). In **A.**, **B.,** and **C., the** widefield maps are overlaid with the results of linear mixed-effect models comparing predicted activity across groups at each timepoint (1 model per ROI). P values were adjusted for multiple comparisons across groups, time and rois by controlling the false discovery rate (Benjamini-Hochberg correction). Blue asterisks indicate significances for the corresponding outlined ROI: *p<0.05, **p<0.01, *** p<0.001.

Correlation analyses further identified that contralesional motor cortex activity was closely related to support-limb performance metrics, including grasp success and accuracy for all groups at late stages of the experiment (late reward rate: Spearman r=0.360, p=0.005; late accuracy: Spearman r=0.561, p<0.001; Figure 4E). Only weak correlations were observed between task-limb driven contralesional activity and task performance metrics (reward rate at the late phase of the experiment: Spearman r=−0.052, p=0.695; late accuracy: Spearman r=0.264, p=0.042; Figure 4D), comparable to those seen for unrelated behavioral variables (reward rate at the late phase of the experiment: Spearman r=0.202, p=0.122; late accuracy: Spearman r=0.285, p=0.027; Figure 4F). Together, these findings indicate that although contralesional cortical activity was enhanced, particularly in animals exhibiting superior motor outcomes following rehabilitative training, this activity did not directly mediate recovery of the impaired limb. Instead, it primarily reflected compensatory postural adjustments of the intact, support limb during skilled grasping behavior.

### Intensive motor rehabilitation increases transcallosal and brainstem axonal sprouting from the contralesional hemisphere

Modeling cortical activity in the ipsi- and contralesional cortex using ridge regression could not sufficiently explain the sensorimotor improvement we measured, in particular in animals with intensive motor training after stroke. Considering the inherent lack of specificity in the cortical widefield signals, we therefore conducted a systematic analysis of structural reorganization—from cortical to subcortical, brainstem, and spinal levels—to elucidate the anatomical substrates underlying the improved motor outcomes in the Rehab group. To assess axonal remodeling, we injected the anterograde tracer Biotinylated Dextran Amines (BDA) into the intact contralesional motor cortex after post-stroke day 28 (Figure 5A), a technique previously established to quantify post-stroke axonal plasticity and structural rewiring^13,17^. Quantification of newly formed transcallosal projections from the contralesional to the ipsilesional hemisphere revealed the greatest increase in fiber density in animals receiving rehabilitative training compared with untreated stroke animals (normalized transcallosal fibers: Rehab group, 0.1725 ± 0.0; Stroke group, 0.078 ± 0.0; Kruskal–Wallis test; Figure 5B). Moreover, motor outcome across all stroke animals at 28 days post-stroke positively correlated with the number of BDA⁺ transcallosal fibers connecting the two hemispheres (Figure 5C).

**Figure 5.**
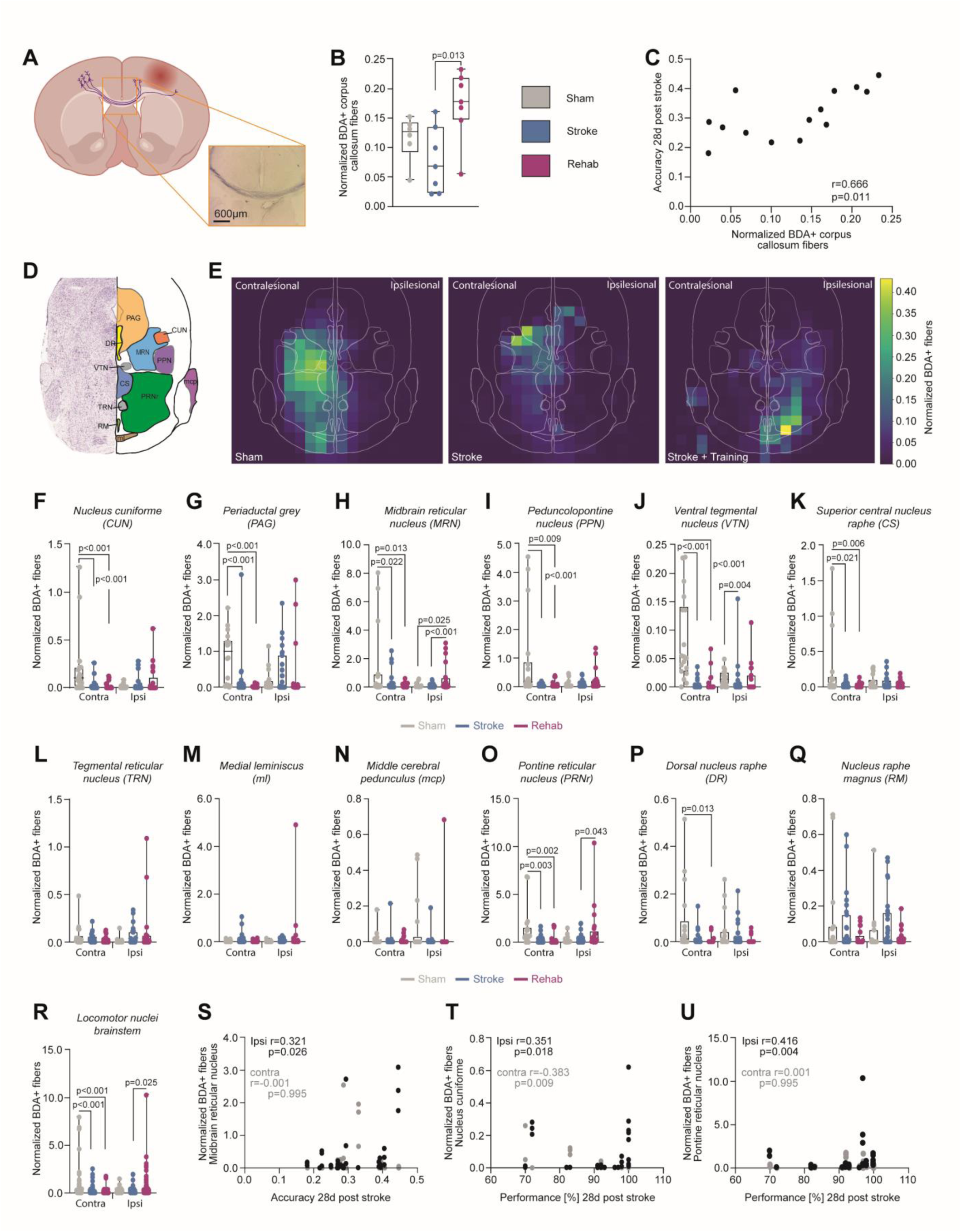
Intensive motor rehabilitation increases transcallosal and brainstem axonal sprouting from the contralesional hemisphere. **A.** Schema depicting biotinylated dextran amine (BDA) tracing of transcallosal fibers from the intact, contralesional to the ipsilesional hemisphere. Inset shows BDA+ transcallosal fibers. **B.** Normalized BDA+ transcollosal fiber counts detected for all experimental groups (Sham n=6, Stroke n=7, Stroke + Training, n=7). **C.** Spearman correlation of BDA+ transcallosal fibers and motor outcome in the lever-pressing tasks for all stroke animals 28 days after stroke. **C.** Scheme depicting brainstem nuclei on a coronal brainstem section aligned to the Allen Mouse Brain Reference Atlas. Abbreviations: PAG = Periaductal grey; DR = Dorsal nucleus raphe; CUN = Nucleus cuniforme; MRN = Midbrain reticular nucleus; PPN= Peduncolopontine nucleus; VTN = Ventral tegmental nucleus; CS = Superior central nucleus raphe; PRNr = Pontine reticular nucleus; TRN = Tegmental reticular nucleus; RM = Nucleus raphe magnus; ml = medial leminiscus; mcp = middle cerebral pedunculus; **D.** Average heatmaps per experimental group displaying normalized fiber densities of BDA+ fibers measured at contra- and ipsilesional brainstem nuclei for the three experimental groups (Sham, Stroke and Stroke + Training group). **F.-Q.** Detailed statistical analysis of distinct brainstem nuclei in 3 brainstem slices per animal per group comparing normalized BDA+ fiber counts between experimental groups on the ipsilesional or contralesion brainstem site. **R.** Statistical analysis comparing BDA positive fiber densities between experimental group in brainstem nuclei dedicated to locomotion including the Nucleus cuniforme, Peduncolopontine nucleus, Pontine reticular nucleus, Midbrain reticular nucleus, Middle cerebral pedunculi and the Tegmental reticular nucleus. **S.** Spearman correlation between normalized BDA+ fibers in the Midbrain reticular nucleus and grasping accuracy in the lever-pressing task 28 days after stroke. **T.** and **U.** Spearman correlations between normalized BDA+ fibers in the Nucleus cuniforme (**T.**) or the Pontine reticular nucleus (**U.**) and the task performance in the grasping task at the experimental endpoint (28 days after insult). Statistical comparison for **B.** and **F.**-**Q.** was performed with a Kruskal-Wallis test setting a significance level of p<0.05.

We next examined whether contralesional projections extended beyond the cortex to subcortical and brainstem targets. While overall normalized fiber density within the basal ganglia and brainstem did not differ significantly among groups (Supplementary Figure 5A-C; Kruskal–Wallis test), brainstem nucleus-specific analyses revealed distinct patterns of axonal sprouting (Figure 5D, E). In sham animals, labeled fibers predominantly targeted contralesional brainstem nuclei. In stroke animals without training, overall fiber counts were reduced, but included sparse projections to ipsilesional brainstem regions. By contrast, animals undergoing intensive rehabilitation exhibited pronounced sprouting into ventrally located ipsilesional nuclei (Figure 5E).

Detailed quantification across individual nuclei (Figure 5D) showed that most contralesional targets—including the nucleus cuneiformis, periaqueductal grey, midbrain and pontine reticular nuclei, pedunculopontine nucleus, ventral tegmental area, and superior central nucleus raphe—displayed either reduced or unchanged BDA⁺ fiber density in stroke groups relative to sham controls (Figures 5F–K, O, P; Kruskal–Wallis test). In contrast, the Rehab group exhibited significant increases in axonal projections to ipsilesional nuclei implicated in locomotion, muscle tone, and posture regulation^20,21^. Specifically, we observed a marked enhancement of fibers targeting the ipsilesional midbrain reticular nucleus (Rehab group, 0.63 ± 0.0; Stroke group, 0.08 ± 0.0; Sham group, 0.09 ± 0.0; Kruskal–Wallis test; Figure 5H) and the ipsilesional pontine reticular nucleus (Rehab group, 1.21 ± 0.0; Stroke group, 0.36 ± 0.0; Kruskal–Wallis test; Figure 5O).

When considering all locomotion-related nuclei (nucleus cuneiformis, pedunculopontine nucleus, pontine and midbrain reticular nuclei, middle cerebellar peduncle, and tegmental reticular nucleus), ipsilesional fiber density was consistently elevated in the Rehab group compared to the untreated stroke animals (Figure 5R). Correlating fiber density with behavioral performance at 28 days post-stroke revealed that axonal sprouting into the ipsilesional midbrain reticular nucleus, nucleus cuneiformis, and pontine reticular nucleus positively correlated with task accuracy and success in the lever-pressing assay (Figures 5S-U; Spearman correlation). In contrast, fiber density in the corresponding contralesional nuclei was uncorrelated or negatively correlated with motor outcomes (Figures. 5S–U). Together, these results suggest that intensive motor training after stroke increased connectivity via transcallosal fibers between the ipsilesional and contralesional hemisphere and drives selective sprouting from the contralesional motor cortex into ipsilesional brainstem nuclei—particularly the midbrain and pontine reticular nuclei— thereby establishing corticofugal detour pathways associated with improved motor recovery.

### Intensive motor rehabilitation prevents unspecific axonal sprouting in the spinal cord

We next examined corticospinal remodeling by assessing BDA⁺ axonal projections from the contralesional motor cortex to the spinal cord. Previous work^13,17^ reported the crossing of axonal fibers of the contralesional, corticospinal tract from the healthy to the stroke-denervated hemi-spinal cord labeled as “midline crossing fibers” (Figure 6A). Quantification of normalized BDA⁺ fiber counts within the stroke-denervated hemicord revealed no significant differences among groups (Sham, Stroke, and Rehab group; Kruskal–Wallis test; Figure 6B), nor in the number of midline-crossing fibers (Figure 6C). However, segment-specific analyses uncovered distinct patterns of axonal sprouting. In untreated stroke animals, BDA⁺ fiber density was increased in the proximal cervical segment C2 (Stroke, 0.77 ± 0.37; Sham, 0.23 ± 0.03; Rehab, 0.14 ± 0.02; two-way ANOVA with Bonferroni post hoc; Figure 6D). In contrast, animals receiving intensive rehabilitative training displayed pronounced BDA⁺ fiber sprouting in the distal cervical segment C8 (Rehab, 0.79 ± 0.38; Stroke, 0.24 ± 0.09; Sham, 0.09 ± 0.01; Figure 6D), suggesting that training promotes reorganization of spinal circuits specifically within distal segments critical for fine motor control.

**Figure 6.**
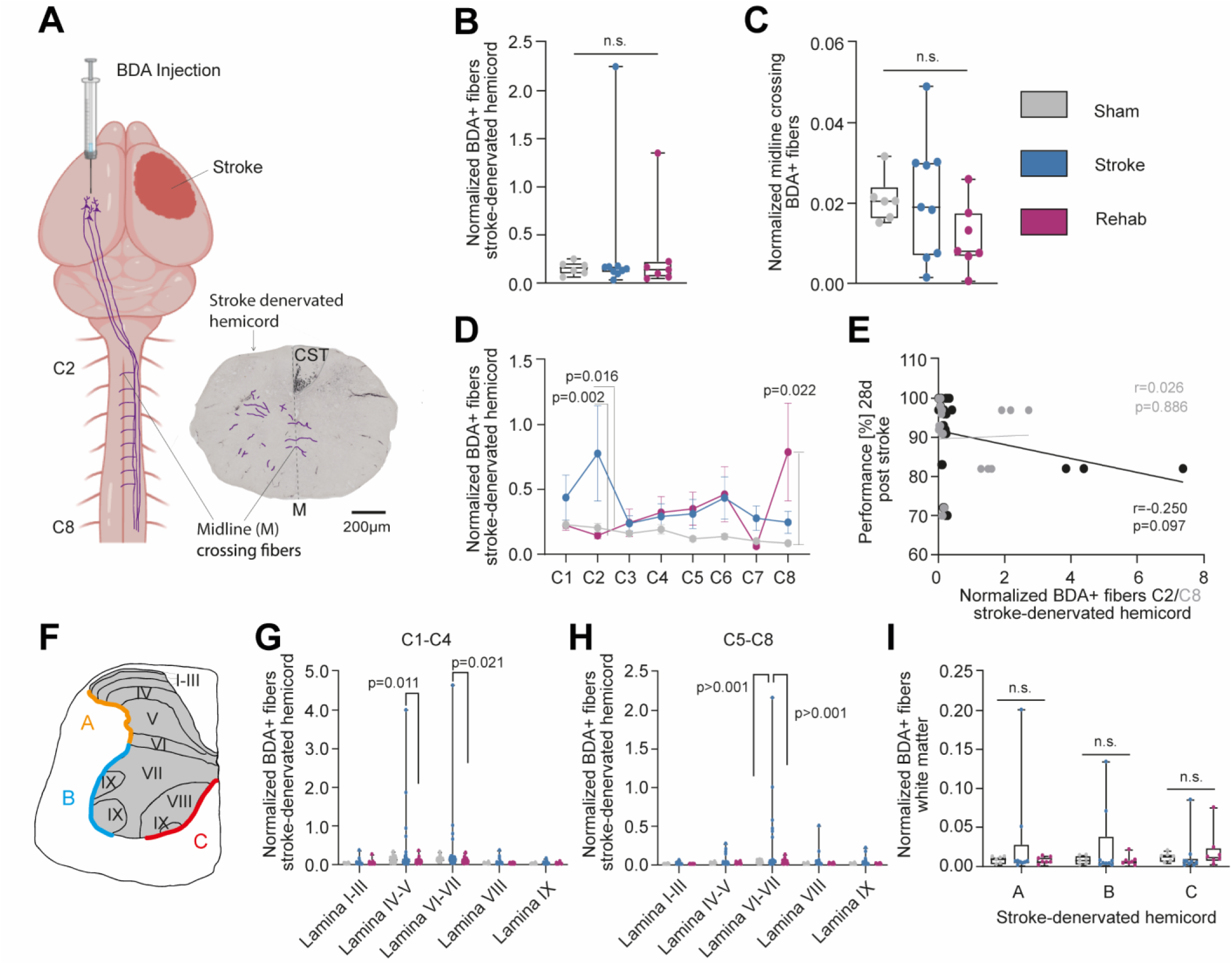
Intensive motor rehabilitation prevents unspecific axonal sprouting in the spinal cord. Scheme displaying BDA-labeled corticospinal (CST) fibers from the contralesional hemisphere, crossing sites twice - at the Decussation pyramidum of the medulla oblongata and at the spinal cord level as midline crossing fibers in the stroke-denervated hemispinal cord (insert, exemplary at spinal cord level C5). **B.** Total amount of BDA+ fibers found in the stroke-denervated hemispinal cord normalized to the amount of BDA+ fibers in the CST in all three experimental groups (Sham n=6, Stroke n=9, Stroke + training group n=7). **C.** Normalized BDA+ fibers directly crossing the midline between both hemispinal cords for all experimental groups. **D.** Normalized BDA+ fibers per cervical segment of the stroke denervated cervical hemispinal cord for all experimental groups. **E.** Pearson correlation between normalized BDA fiber count either at cervical level C2 or C8 and the motor performance in the lever-pressing task at final endpoint (28 days after stroke). **F.** Exemplary scheme depicting the laminae (I-IX) in the grey matter of the spinal cord at cervical level C5 as well as the gray matter-white matter boundaries in the dorsolateral (labeled with “A”), the ventrolateral (label “B”), and the ventro-medial funiculus (label “C”). **G.** and **H.** Lamina-specific analysis of BDA+ fibers in the stroke-denervated proximal (C1-C4, **G.**) and distal (C5-C8, **H.**) cervical hemispinal cord for the three experimental groups. **I.** Normalized BDA+ fibers crossing into the white matter at A, B and C. For **B.**, **C.**, statistical comparison was performed with a Kruskal-Wallis test. For **G.-I.** a two-way ANOVA with posthoc Bonferroni was executed. The significance level was set at p<0.05.

We next examined whether the extent of axonal sprouting within the denervated hemicord correlated with behavioral recovery 28 days post-stroke. Neither fiber density in segment C2 nor C8 was correlated with task performance in the lever-pressing assay (Pearson’s correlation; Figure 6E). To elucidate this effect, we analyzed fiber sprouting in the different laminae of the gray matter (Figure 6G) at proximal (Figure 6H) and distal (Figure 6H) cervical segments. In untreated stroke animals, sprouting was prominent within laminae IV–V, corresponding to proprioceptive regions, and laminae VI–VII, associated with sympathetic control (Figure 6H). In contrast, neither stroke group exhibited enhanced targeting of motor neuron pools in lamina IX (Fig. 6G, H). Excessive sprouting at gray– white matter boundaries within the dorsolateral, ventrolateral, or ventromedial funiculi was not observed in any group (Figure 6I). Although we found structural rewiring in the spinal cord with specifically enhanced fiber sprouting into the stroke denervated-spinal hemi cord at cervical segment C8 for animals with intense rehabilitative training, this sprouting was not directly linked to improved outcome. But intense training prevented maladaptive, nonspecific sprouting in dorsal and medial gray matter, as seen in stroke animals without further training.

## Discussion

We demonstrated that large sensorimotor strokes induce chronic motor deficits, which can be only partially rescued by intense rehabilitation strategies (Figure 1). Chronic wide-field calcium imaging across both hemispheres - encompassing the ipsi- and contralesional cortices - during acquisition of a skilled lever-pressing task and over several post-stroke weeks enabled longitudinal mapping of cortical reorganization associated with motor recovery. Intense motor training enhanced contralesional cortical activity and coincided with improved behavioral recovery; however, this increased activity was not directly linked to fine motor performance of the affected limb (Fig. 2). Ridge-regression modeling indicated that contralesional activation predominantly reflected movements of the intact support limb and contributed only marginally to the restoration of impaired-limb function (Figs. 3, 4). Using systematic tracing studies, with stereological investigation of axonal sprouting from the contralesional motor cortex to subcortical and spinal cord areas, revealed anatomical correlates in the form of corticofugal circuit rewiring to distinct ipsilesional brainstem nuclei as critical components related to improved motor outcome after stroke (Figure 5).

Our study contributes to the ongoing debate concerning the role of the contralesional cortex in functional recovery following large strokes. Several preclinical and clinical studies have shown increased cortical activity in the contralesional hemisphere^2,5,8,22–24^. However, interpreting these activity patterns in relation to functional outcome has proven challenging, yielding seemingly contradictory findings: both stimulation and inhibition of the contralesional cortex have been reported to enhance post-stroke recovery^25–30^. Our study underscores that causal inferences drawn from correlative observations remain inherently problematic. For example, in our study increased contralesional activity in the motor cortex correlated with improved functional performance, exemplified by a greater number of successfully executed grasps (Fig. 2F). However, a detailed analysis of motor behavior including fine skilled motor features of the impaired limb demonstrated no positive effect (Figure 2F), indicative that a sophisticated in-depth analysis of outcome parameters is essential to make valid claims connecting activity patterns to functional recovery mechanisms.

Furthermore, the interpretation of cortical activity patterns requires the consideration of the full posture of the animal including uninstructed movements or task specific motor responses such as licking behavior during the reward^31^. By applying a ridge regression model that incorporated task-related movement parameters—including support-limb motion, responses to auditory cues, and licking behavior during reward—we were able to disentangle the distinct behavioral components driving cortical activity across regions (Figs. 3, 4). This analysis provided mechanistic insight into the ongoing debate regarding the role of the contralesional cortex, revealing that enhanced contralesional activity primarily reflects movements of the supporting limb rather than genuine recovery of fine motor features in the stroke-impaired limb. In particular, we found enhanced contralesional activity in animals, which had received intense motor training after stroke. This rehabilitative strategy failed to achieve full motor restitution (*restitutio ad integrum*), as fine motor features such as hand aperture and finger flexion remained chronically impaired (Fig. 1H). Thus, the improvement of task performance, success in rewarded grasps and accuracy (Figure 1E, F, G) can be explained in the context of compensational strategies, including adapted posturing of the support limb but also enhanced coordination between the support limb and the impaired task limb– through improved connectivity between both hemispheres (Figure 2, 4, 5). Our findings indicate that focusing solely on single outcome parameters while neglecting compensatory strategies obscures the complexity of structural and functional reorganization after stroke. A broader consideration of compensatory movements and postural adjustments is essential to avoid misleading interpretations and erroneous causal inferences.

Our systematic top-down analysis of BDA tracings from the contralesional motor cortex, aimed at identifying anatomical correlates underlying the improved outcomes observed after intensive rehabilitative training, revealed structural reorganization across multiple anatomical levels. In animals of the Rehab group, we found pronounced transcallosal axonal sprouting connecting the ipsi- and contralesional hemisphere. This enhanced transcallosal connectivity was positively correlated to improved motor outcome parameters (Figure 5B, C). Our results are in accordance with findings from several other studies^32,33^ and suggest an improved communication between both hemispheres, facilitating the coordination of postural adjustments and compensatory movements of both, the task and the support limb. However, our study goes beyond and identifies a detour pathway from the contralesional motor cortex, targeting specific ipsilesional brainstem nuclei such as the midbrain and pontine reticular nucleus (Figure 5H, O) and relating the increased fiber density in these ipsilesional nuclei to motor outcome parameters at the endpoint of our study (Figure 5S-U).

Several studies have indicated the importance of brainstem reorganization for the recovery of motor function after stroke^20,34–36^. However, here we demonstrated the specific targeting of the ipsilesional midbrain and pontine reticular nucleus as a specific corticofugal pathway induced by intensive rehabilitative training connecting the contralesional motor cortex to ipsilesional brainstem nuclei relevant for locomotion, muscle tone, and posture regulation^20,21^. Furthermore, our study identified these motor nuclei of the brainstem as critical relay hubs for structural reorganization, enabling forms of motor recovery and compensatory strategies. Our results suggest that cortical and corticofugal reorganization are the main drivers explaining functional outcome parameters, while reorganization on the spinal cord level may have a rather refining character. We found that enhanced corticospinal sprouting crossing at the cervical spinal cord level from the healthy to the stroke-denervated side (“midline crossing fibers”) was negatively correlated to functional outcome parameters (Figure 6E). We could identify pronounced targeting of midline crossing fibers at distal segments such as C8 (Figure 6D), thus targeting a spinal cord level for fine muscle innervation of hand function rather than proximal innervation of shoulder muscles (as seen for stroke animals without rehabilitative training, Figure 6D). Thus, our data suggest that intense training rather supports the refinement of targeting and maintains mechanisms of pruning, preventing over-excessive sprouting to dorsal and intermediate gray matter parts. In our data, untreated stroke animals showed prominent axonal sprouting within laminae IV–V, corresponding to proprioceptive regions, and laminae VI–VII, associated with sympathetic control (Figure 6G, H), which might have induced installation of interfering, maladaptive spinal cord microcircuits^17,37^, an effect which was absent in animals with intense rehabilitative motor training.

Overall, our findings highlight that recovery of skilled motor function such as goal-directed grasping behavior requires structural and functional reorganization on several levels. We have identified enhanced cortico-cortical reorganization through increased transcallosal connectivity fostering compensatory strategies involving movement adjustments of the healthy support limb and an improved coordination between both paws, the impaired and the healthy one. Furthermore, we identified a rehabilitation-induced corticofugal pathway, from the cortex to distinct brainstem nuclei, which is critical for enhanced motor recovery. Finally, our study also reveals a third level of reorganization on the spinal cord level with further refinement in the structural reorganization of the spinal cord grey matter through rehabilitation, elucidating coordinated targets for neuromodulation strategies in chronic post-stroke impairment.

## Acknowledgements

We thank Martin Wieckhorst for technical advice and fruitful discussions. We thank Beate Aschauer and Astrid Baltruschad for their work and technical support processing the histological data. We thank Philipp Bethge for providing transgenic mice used in this study. We thank Antonia Weingart for helping with figure design and illustrations. This study was supported by the Demenz Forschung/ Synpasis foundation Switzerland (to ASW) and the Swiss National Science Foundation Grant #192678 (to ASW) as well as the TRR 274 – Checkpoints of Central Nervous System Recovery by the DFG awarded to A.S.W.

## Autor contributions

A.S.W. ideated and provided the concept. M.P. and A.S.W. designed the study. M.P. and A.S.W. performed surgeries and carried out experiments. N.T., C.H., and A.S.W. performed the histological analysis. M.P., C.H. and A.S.W. performed the data analysis with input from S.M. and F.H. N.B. provided support for the stereological analysis. U.S. performed MRIs. F.H. provided resources. M.P. and A.S.W prepared figures and wrote the manuscript with input from all authors.

## Supplemental information

Document S1. Table S1 and S2. Figures S1–S5. Supplementary Video 1.

## Methods

### Animals

We used 38 adult mice aged 3 to 6 months at the first day of experiments of both sexes (n=21 males, n= 17 females). Mice were GP5.17 Thy1-CaMP6f +/− transgenic mice (Jackson Laboratory, RRID: IMSR\_ JAX:025393^38^. 17 animals were excluded because of complications of the stroke surgery (n=5), too severe motor deficits (n=3) or no deficit at all (n=1), because animals did not learn the grasping task (n=4) or the imaging and tissue quality was not sufficient enough for further processing (n=4). Mice were housed in groups of two to four in standard cages (530 cm² floor area, 7.4 L) under controlled conditions: a 12-hour light/dark cycle, a constant temperature of 22 ± 1 °C, and free access to food and water. All experiments were conducted during the animals’ active (dark) phase and in compliance with the guidelines of the Swiss Federal Veterinary Office, under license ZH086/2022 approved by the Cantonal Veterinary Office in Zurich. The study design also adhered to the Stroke Therapy Academic Industry Roundtable (STAIR) recommendations^39^ for preclinical stroke research. Sample sizes for each experimental group were determined based on the means and variances reported in related studies^13,17,34,40^, ensuring sufficient power (p < 0.05, power > 0.8) to detect statistically significant effects in ANOVA analyses.

### Experimental set-up and task

The behavioral setup was adapted from Mathis et al,^41^ and consisted of a 2-axis joystick sensor (M7 Hall Sensor Gimball, FrSky) and thin carbon rod, attached to a custom-machined aluminium holder. The springs in the base of the joystick were adjusted so that a 1mm deflection would require less than 0.1N of force. The mice were placed in a head fixation stage and were provided with a Q-Tip for paw support. The joystick was placed at 11 mm distance from the paw support, centrally with respect to the paw rest positions, and the tip was at a 5 mm elevation relative to the paw rest. Deflections were measured as a voltage from the hall-sensors inside the joystick, digitized via a NI DAQ card (NI USB-6001, National Instruments), and recorded on a computer via a custom LabView program. A servo motor (DES 718 BB MG, Graupner) was used to deflect the joystick out of reach of the mouse during intertrial periods.

Two high-speed behavioral cameras (acA1300-200um, Basler) with 12.5 mm focal length objectives (HF12.5HA-1S, Fujifilm), were placed symmetrically at either side of the joystick, and recorded videos of each forepaw at 150 Hz (1280×1024 pixel resolution). The camera recordings were synchronized, compressed, and saved using custom LabView code.

Auditory cues were provided by a pair of speakers (Z150, Logitech) and consisted of 3 x 50 ms pure tone beeps at 8 kHz, separated by 50 ms. Sweetened water droplets (approx. 1uL, 10g sugar in 200ml water), were provided as rewards via a metal spout, and controlled by a solenoid valve (VDW22JA, SMC Pneumatics).

The behavioral task was set up to assess reach-to-grasp behavior. Head-fixed mice were trained to reach and pull the joystick towards themselves. Upon a successful deflection (10mm), mice received a reward. Each trial started with a 2 s pre-trial period, followed by the auditory cue. The joystick was moved into place using the servo motor while the cue played. If mice pulled the joystick successfully within a 5 s window, they immediately received a water reward, and an intertrial period of 4 s started. During the intertrial period the joystick was removed from reach by the servo motor. To ensure the mice stayed motivated during long bouts of unsuccessful trials, there was a 10% chance of receiving a reward after an unsuccessful trial. To avoid over-training the task, each experimental session consisted of exactly 100 trials.

### Behavioral training

Following the imaging preparation surgery, mice were extensively habituated to handling, and then to tolerate head fixation of increasing intervals for a period of 2 weeks. To ensure motivation in the task, mice were placed on a water scheduling regime by adding citric acid (2% solution, Sigma-Aldrich) to their in-cage drinking water^42^. The water scheduling remained in place for the entire duration of the experiment. Once per week, mice had ad-libitum access for 1h to un-soured water, and their weight was monitored consistently to ensure that it did not fall below 15% of the starting weight. The water scheduling was interrupted 1 day prior to photothrombotic stroke surgery and resumed 2 days after the surgery.

Once habituated to head-fixation, mice were trained for 2 days to lick the water delivery spout, receiving rewards at random intervals with 6 second average spacing. Following this, mice were trained to touch the joystick, by rewarding any form of contact between the mouse and the joystick. Once the mice were consistently touching the joystick (>1 mL rewards received in a session), mice were rewarded for small (approx. 1mm) deflections of the joystick. The size of deflection necessary to elicit a reward was gradually increased until mice were consistently (1 mL rewards in a session) pulling the joystick for 1mm towards themselves. This process took 5-8 sessions on average.

Finally, mice were introduced to the full task structure and were trained daily in 100-trial sessions, while simultaneously recording neuronal activity using widefield calcium imaging. Mice reached expert performance in the task (>80% successful trials for 3 consecutive sessions) on average after 10 +/− 0.4 sessions.

Mice were free to perform the task with either forelimb. A clear limb preference typically emerged during the first attempts to reach the joystick, and remained stable throughout the experiment, including in the presence of stroke-induced limb impairments. The limb used to perform the task was defined as the “task limb”, while the other was defined as “support limb”. The hand preference distribution (n=11 left-handed, n=10 right-handed) is consistent with known laterality of naïve mice from C57BL/6 backgrounds^43^.

### Experimental timeline and cohorts

All different experiments were repeated in N=4 independent studies with n=4-10 animals per study for behavioral and neuronal analysis. As there was no statistically significant difference in the outcome of lesions, behavior, and anatomy, the data shown here were pooled from all studies. The final histological analysis was performed by two independent investigators who were not involved in training, testing, and imaging the animals. This study conforms with the AARIVE guidelines (https://www.nc3rs.org.uk/arrive-guidelines). After reaching expert performance in the task (>80% successful trials for 3 consecutive sessions), mice were randomly assigned to an experimental cohort. As stated above, the number of animals included in this study was n=21, distributed as follows. “Stroke” (n=9) and “Rehab” (n=7) mice received a photothrombotic stroke, while “Sham” (n=5) received the sham photothrombotic intervention. Post-stroke, all experimental cohorts performed the task during widefield imaging at fixed timepoints: post-stroke days 3, 7, 14, 21 and 28. For the purposes of data analysis, days 3-7 are referred to as “Early” post-stroke and days (14,21,28) are referred to as “Late” post-stroke sessions.

Mice belonging to the “Rehab” cohort performed 4 additional behavioral sessions per week (100 trials each), starting on day 8 post-stroke. Comparisons between all groups were performed on the original day 3,7,14,21,28 sessions.

### Surgeries

For all surgical procedures, mice were deeply anesthetized with 4% isoflurane delivered in 700–800 mL/min O₂. Buprenorphine (0.1mg/kg bodyweight, Temgesic®, Reckitt & Benckiser) was injected intraperitoneally 30 minutes prior to the surgery. Vitamin A cream (Bausch & Lomb) was applied to both eyes, and body temperature was maintained at 36.5 °C using a heating pad. During surgery, animals were secured in a stereotaxic frame (Kopf Instruments) and maintained under 1.5-2% isoflurane anesthesia. Following surgery, mice were kept on a heating pad until fully recovered and ambulatory. Postoperative analgesia was provided with Buprehorphine (0.1mg/kg bodyweight, 4h and morning post-surgery) and subcutaneous Meloxicam (5mg/kg, Metacam®, Boehringer Ingelheim) every 12 h for 1 day, followed by daily administration for an additional 2 days. For the non-invasive photothrombotic stroke procedure, no analgesics were administered.

#### Preparation for chronic wide field calcium imaging in vivo

Mice were fixed in a stereotaxic frame under anesthesia as described above. The scalp was shaved, sterilized with a Betadine solution, and Lidocaine crème was applied for local analgesia. A circular patch of skin was cut to reveal the skull, and any connective tissue was cleaned from the surface using cotton swabs and saline solution. A circular ring of UV-curable dental cement (Charisma®, Kuzler) was applied around the edge of the skull, and a titanium plate for head-fixation was fixed to the posterior part of the preparation using additional dental cement. The edge of the preparation was coated with a layer of black self-curing dental acrylic (Ortho-Jet, Lang Dental) to minimize autofluorescence of the Charisma acrylic during imaging. The surface of the skull was coated with 4-5 thin layers of cyanoacrylate glue (Pacer Technologies), with 1-2 minute pauses to let each layer dry. Finally, tissue glue (Vetbond, 3M) was used to fix any loose skin in place.

#### Photothrombotic stroke

To induce ischemic strokes, the animal was stereotaxically fixed as described above. The skull was covered with an opaque template (15×15mm, with an opening of 5×3mm) and aluminum foil to protect other regions of the brain and the eyes except for the region to be lesioned. We then injected (i.p.) 0.1ml of a freshly prepared light sensitive dye (10 mg/ml Rose Bengal in 0.9% NaCl solution, Sigma-Aldrich). After a waiting period of 8-10 min, the skull of the sensorimotor cortex was exposed to a strong light source (Olympus KL 1500LCS, 150 W, 3000 K) for 8-10 min inducing a photothrombotic stroke destroying the sensorimotor cortex of the preferred forelimb. For left-handed mice, this was the right sensorimotor cortex (A/P −2 to +3mm, M/L 0 to + 3mm) and for right-handed mice, this was the left sensorimotor cortex (A/P −2 to +3mm, M/L −3 to 0mm). Sham-operated mice underwent identical surgery, without turning on the light source during the illumination period. In the main text, we identify the dominant-forelimb sensorimotor cortex as “ipsilesional”, and the non-lesioned cortex as “contralesional”.

#### BDA injections into the intact motor cortex

Anterograde tracing of structural rewiring from the intact motor cortex was performed 5-6 weeks after stroke using Biotinylated Dextran Amine (BDA, 10,000 molecular weight, 10% solution in 0.01 M PBS, Invitrogen) as previously described^13,17^. For this procedure, animals were put in isoflurane anesthesia and placed in a stereotaxic frame as described above. We injected BDA at four injection sites (200 nL each) in the intact contralesional motor cortex.

| Location<sup>44</sup> | A/P | M/L (right hemisphere) | Depth |
| --- | --- | --- | --- |
| Forelimb Area I | +2.2 mm | +0.95 mm | -0.6 mm |
| Forelimb Area II | 0.0 mm | +1.75 mm | -0.5 mm |
| Hindlimb Area I | -1.25 mm | +0.88 mm | -0.6 mm |
| Hindlimb Area II | -1.5 mm | +1.75 mm | -0.5 mm |

### Analysis of motor behavior

The video recordings for each paw were spatially downsampled by a factor of 1.5 and processed using DeepLabCut^18^, to extract the following keypoints for each forelimb: elbow, wrist and 3 keypoints per finger (knuckle, middle and tip). Keypoints with a low posterior probability value (< 0.5) were removed and replaced via linear interpolation. Finally, the time series of each keypoint coordinate were filtered with a second order Savitzky-Golay filter with a window size of 75 ms.

The behavioral data was segmented into “grasps” via a thresholding-based approach. First, for each forelimb, the forelimb position time-series was computed as the average position across all tracked keypoints, and the absolute velocity of the paw was computed by taking the L2 norm of the x,y coordinate temporal derivatives. The velocity trace was then thresholded to yield the onsets and offsets of movements. A task-limb movement was considered “rewarded”, if the onset immediately preceded a successful deflection of the joystick. “Missed” attempts were classified as movements whose trajectories crossed the location of the joystick on the camera, but did not result in a reward, and all other movements were classified as “other”.

Grasping accuracy was defined as the percentage of rewarded grasps to rewarded plus missed grasps. Task performance was quantified as the percentage of rewarded trials in a session. To account for differences in trial length, we compute a “reward rate”, which is the task performance divided by the duration of the session in seconds.

The coordinates of the tracked paws were used to extract features, representing motor kinematics of interest. The limb rotation was defined as the mean angle across fingers with respect to the vertical and reflects proximally driven rotations of the entire forepaw. The hand aperture was defined as the angle between the first and last finger and the finger bending was defined as the mean, across fingers, of the angle between the initial and final phalanx of each finger (angle between knuckle-to-middle and middle-to-tip segments).

For analysis where the behavioral data was directly compared to the widefield imaging data, the behavioral timeseries were downsampled to the widefield sampling rate via linear interpolation.

#### Matched movements analysis

To assess whether changes in cortical activity were dependent on the specific trajectory of the movements (Supplementary Figure 3), we devised a procedure to reduce the variability within the set of rewarded grasps. First, for each rewarded grasp we computed a set of continuous features in a −50 to 300ms window (53 frames at 150Hz sampling rate) around the grasp onset. These features were the distance to the joystick, the velocity of the limb, the limb rotation, hand aperture, and finger bending, resulting in a (5, 53) matrix for each grasp.

We then computed a pre-stroke grasp template for each mouse, by taking the average over all pre-stroke grasps. For each grasp in the dataset, we computed its shape-based distance (SBD)^45^, to the template grasp. For each mouse, we computed a distance threshold, based on the median SBD of all pre-stroke grasps to their respective template. Any grasps with SBD below this threshold were considered similar to the pre-stroke template and included in subsequent analysis.

We additionally computed the same features, for the support limb, still aligned to rewarded grasps only. We repeated the same analysis, resulting in a set of rewarded task-limb grasps which shared similar movements of the support limb during the grasp execution.

### Widefield calcium in-vivo

Widefield calcium imaging of neuronal population activity was performed through the intact skull using a custom-built widefield microscope, with a dual front-to-front objective design^46^. Two excitation LEDs with central wavelengths of 470nm and 405nm (M470L3 and M405L4, Thorlabs) were used for GCaMP excitation and hemodynamics correction respectively. Both LEDs were filtered using bandpass filters of 10nm width around the LED’s central wavelength, and the two illumination paths were merged via a dichroic mirror with longpass wavelength of 435nm. The LEDs were synchronized to illuminate alternate frames using custom software on an Arduino microcontroller (Uno Rev3, Arduino). Light from the sample was collected through the objective (D0-5095, Navitar), and re-focused onto a CMOS camera (Orca-Flash 4.0 v3, C13440-20CU, Hamamatsu Photonics), via a second, identical objective, yielding a total magnification factor of 1 and a field of view of 13.5×13.5mm. Emitted fluorescence was additionally filtered with a 525/24nm bandpass filter. Data was acquired at 512×512 pixel resolution, and a sampling rate of 40Hz, resulting in 20Hz per channel.

### Analysis of chronic widefield calcium imaging

The raw imaging data was first spatially filtered with a 5×5 pixel gaussian kernel and spatially binned to 128×128 pixels. Each imaging session was spatially registered to a reference session, using an affine transformation. Slowly varying temporal effects were removed using a high-pass 2^nd^ order Butterworth filter (cutoff frequency: 0.1 Hz), applied to each pixel and channel independently. Each pixel time series was further temporally filtered with a 1 frame gaussian kernel. For each session, a binary mask was hand-drawn, to select the pixels of the imaging field of view belonging to the brain.

To estimate signals representing movement-related artifacts, while preserving signals related to true movement-related neuronal activity, we computed the mean fluorescence timeseries of the 470 nm channel, across all pixels not belonging to the brain (*F*_*movement*_). For each pixel *i*, the scale of the 405nm channel fluorescence 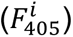 and *F*_*movement*_were matched to the scale of the 470 nm channel timeseries 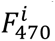, via the following linear regression model:

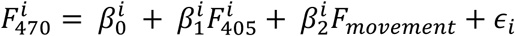

The hemodynamics and movement-corrected fluorescence 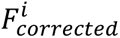 for each pixel was defined as the residuals of the fit, using the estimated parameters *β̂*^*i*^:

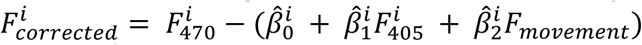

For each pixel, the (Δ*F*/*F*_0_)^*i*^ was calculated by dividing the mean fluorescence across time (over the entire session) for each pixel, 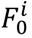:

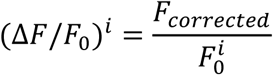

Finally, for analysis of movement-aligned, we subtracted the mean activity over periods where the mouse was completely still, so that the zero-point of activity was relative to quiet periods. For computational and storage efficiency, the was compressed using Singular Value Decomposition (SVD), keeping the top 100 components, which captured > 99% of the variance in each session. All analyses were performed in SVD space and transformed back into pixel space for reporting results.

When averaging across mice, all sessions were first spatially registered using an affine registration to the Allen Atlas Common Coordinate Framework^19,46^, based on the following anatomical landmarks: bregma, lambda, the medial base of the retrosplenial cortex, and the frontal meeting point of the cortex with the olfactory bulb. The widefield maps for right-handed mice were mirrored along the x-axis, so that the ipsilesional hemisphere is always shown on the right, and the contralesional hemisphere is shown on the left in all figures. We extracted time courses for each Allen Atlas region of interest (ROI) by averaging the of all pixels contained within a given ROI. For statistical comparisons across, we defined widefield “responses” as the average over a 1 s window following the onset of a rewarded grasp.

### Ridge Regression Models

To investigate the contribution of different behavioral variables to cortical activity, we set up ridge-regression models for each experimental session^31^. We set it up as a multiple linear regression model, with the target variable being the SVD-compressed cortical activity matrix of shape (*N*_*frames*_, 100 components). As regressors, we used binarized variables, which were set to one at frames representing onsets of the following behavioral events: task-limb movement onsets, support-limb onsets, auditory cue and reward delivery. We additionally included time-lagged versions of each regressor. For task-limb and support limb movements, we included lags from −3 to +3 seconds, in single-frame increments, while for “task-variables” (cue + reward), we included increments from 0 to +3 seconds.

The models were fitted using the scikit-learn library for Python 3.10^47^. The ridge regularization strength was estimated via 5-fold cross-validation. To score the models, we generated the cross-validated model predictions and computed the *R*^2^ in pixel space. We additionally generated (cross-validated) partitioned model predictions by multiplying subsets of the design matrix by subsets of the estimated coefficients representing each behavioral variable.

As an additional measure of model performance, we computed the unique explained variance Δ*R*^2^ of each behavioral variable, by creating reduced models where the variable of interest (+ time-lags) was randomly permuted along the time axis. The Δ*R*^2^ for the variable was defined as the difference in *R*^2^ between the full model and reduced model.

### Histology

After conclusion of all experiments (6 – 8 weeks after stroke), mice were deeply anesthetized with 5% Isoflurane and overdosed with pentobarbital (Kantonsapotheke Zurich, 300 mg/kg body weight, i.p. injection). As soon as respiratory arrest occurred, 0.05 mL Heparin (Braun) was injected into the left ventricle, and the animal was perfused transcardially with cold 0.1M PO4 followed by 4% paraformaldehyde (PFA) in 0.1M PO_4_. Brains and spinal cords were extracted, post-fixed (4% PFA, 4 °C, 24 h), cryoprotected (30% sucrose, 0.1M PO_4_, 4 °C, 48 h), embedded in OCT (Tissue-Tek, Sakura), frozen at −80 °C, and 40 µm coronal sections were cut with a sliding cryostat (Leica). An on-slide staining using the nickel-enhanced DAB (3,3′-diaminobenzidine) protocol (Vectastain ABC Elite Kit, Vector Laboratories; 1:100 in Tris-buffered saline plus TritonTM X-100) followed as previously described^13,17^.

#### Analysis of BDA+ transcallosal fibers

We performed the analysis of all histological slices single blinded. A total of 22 brains were analyzed quantifying axonal fibers in the corpus callosum using a Zeiss light microscope (Axioskop). The corpus callosum was identified in individual brain sections starting in the forebrain to the end of the corpus callosum (beginning of the hippocampus) using a 10x objective. We selected every fourth section from the beginning to the end of the corpus callosum for analysis. Fiber quantification was performed with 20x and 40x objectives using a grid counting the number of BDA+ fibers crossing at the midline between both hemispheres. For normalization, BDA+ fibers were counted in a defined area of the intact and well-stained corticospinal tract at the level of the brainstem. The number of fibers was determined within a standardized region of 0.05 x 2.5 mm². The ratio of BDA+ fibers at distinct areas (corpus callosum, basal ganglia, brainstem, and spinal cord) relative to the BDA+ CST fibers at brainstem level was calculated for further analysis and statistics with group specific comparison.

#### Analysis of midline-crossing BDA+ fibers in the spinal cord

For all coronal spinal cord slices we first performed an assessment of the cervical segment using the spinal cord histological atlas by Anderson et al.,^48^ determining spinal cord segments C1 to C8 using a light microscope (Olympus BX50) with a stereo-investigator system (Stereoinvestigator, Version 2022, MBF Bioscience). To ensure precise anatomical identification of landmarks of the different segments, a dark-field microscope was additionally employed. We counted BDA+ fibers in the stroke denervated cervical hemi-spinal cord with 3-4 sections per segment analyzing a total or approximately 530 spinal cord sections. Quantification of BDA+ fibers was performed using the mode “Systematic Random Sampling (SRS)” of the stereo-investigator software. This counting method ensures that all regions of the structure are sampled with equal probability, while simultaneously providing a systematic and uniform distribution of sampling sites, thus improving representativeness and reducing bias. We defined also counting rules: only fibers inside distinct boundaries of the counting grid were counted. Only clearly identifiable axons with boutons were considered. We contoured the grey matter and generated a counting grid (Perimetrics modul of the stereo-investigator) in the region of interest in the stroke denervated hemi-spinal cord using the following parameters: Counting frame size: 100 × 100 µm, SRS grid size: 150 x 150 µm, Merz radius: 50 µm. For counting, we used a 40x objective and marked all fibers on images taken from the slides while counting with an integrated camera of the Olympus BX50 stereo-investigator system.

To capture midline crossing fibers of the spinal cord, we drew a square at the midline of the spinal cord sections through the central canal. BDA+ fibers crossing this midline were counted as “midline crossing fibers”. The alignment was performed individually for each section, allowing precise morphological allocation.

For the analysis of lamina-specific sprouting we took the images generated with the stereo-investigator (as described above) and overlaid these spinal cord sections with a template showing the boundaries of the different laminae: Laminae 1-3, Laminae 4-5, Laminae 6-7, Laminae 8-9 in the grey matter and areas A, B and C in the white matter (see Figure 5F-I).

#### Analysis of BDA+ fibers in the brainstem

Fibers throughout the brainstem were quantified similarly to those in the spinal cord using the SRS method and stereo-investigator software from MBF Bioscience for unbiased stereological analysis of histological images. Per animal we analyzed 3 brainstem sections. The following parameters were applied in the Stereoinvestigator application “Petricmetrics”: Counting frame 300 x 300 µm, Merz radius: 50 µmm Slides analyzed: 200-350 per brain. A midline was drawn to differentiate the ipsi- and contralesional part of the brainstem. The coordinates of the counting grid including fiber coordinates were saved and compiled into a table and analyzed using a custom python script. For heatmap generation we aligned grids with fiber counts of individual animals in such a way that ipsi- and contralesional parts overlaid. Next, we laid a uniform grid (cell size matching the StereoInvestigator grid over each slice and counted how many fibers fell into each grid square. Counts were normalized by taking the ration of BDA+ fiber counts in the brainstem to BDA+ fiber counts in the CST (as described above). We created heatmaps of fiber counts in the brainstem by averaging the results of 3 brainstem sections per animal and finally averaged across animals of a treatment group (“Sham”, “Stroke”, “Stroke plus training”).

For the analysis of BDA+ fibers in different brainstem nuclei we took images of the brainstem sections with the overlaying counting grid of the StereoInvestigator progam and used the Allen mouse brain atlas^19^ to identify individual brainstem (Figure 5D) nuclei. We delineated those on the brainstem sections including our counting grid and applied a custom python script allocating the amount of BDA+ fibers in the counting grid to distinct brainstem nuclei.

### MRI

To examine stroke lesion size and location PFA-fixed brains were scanned ex-vivo by a 7T small animal MR system (Bruker BioSpin GmbH, Ettlingen, Germany). Prior to imaging brain samples were placed in 15 ml falcon tubes filled with perfluoropolyether (Fomblin®Y-LC 80, Solvay Solexis, Bollate, Italy). The brain MR measurements were accomplished using a volume resonator for excitation and a four-element phased array surface coil for signal detection. 48 coronary slices of 0.32 mm thickness resulted in T2-weighted images which were acquired using a TurboRARE sequence with the following parameter: matrix dimension= 200 x 200, spatial resolution= 50um x 50um voxels, mean diffusivity= 200 x 140,repetition time= 350 s, echo time=10um, number of averages= 120. After ex vivo MRI, brains were removed from Fomblin and again placed in PFA. The quantification of the lesion volume was performed with Image J, counting the pixels in the encircled area of interest. We also extrapolated the pixel count for the stroke volume to the corresponding volume of the unaffected contralesional hemisphere, taking the shrinkage of stroke scar tissue into account.

### Data management and statistical analysis

Experimental data and analysis pipelines were managed by a custom DataJoint database^49^ (RRID:SCR\_014543) implemented in Python v3.10. Statistical analysis and plotting were performed with R v4.5.1, Python and Prism v10 (Graphpad), while figures were assembled with Adobe Illustrator (v28.3).

Unless stated otherwise, data are reported as mean ± standard error of the mean (SEM). Data points in figures represent all measured samples (e.g. individual grasps or trials) unless stated otherwise. To estimate changes in a given response variable between experimental groups during different stroke phases, we implemented linear mixed-effect models. The response variable was modeled as linear combination of two categorical factors (including interactions): the stroke group and the experimental phase. To account for repeated measures within mice and days (e.g. multiple grasps per mouse at a given day), we implemented random effects per unique mouse identifier and experimental day. Such models were specified in R as follows:

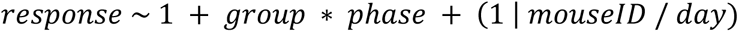

The models were fit using the lme4 (v1.1-37) package and pairwise comparisons between groups were computed on the model estimated marginal means^50^ using the emmeans (v1.11.1) package^51,52^. When modelling the responses of multiple imaging ROIs, we fit one model per ROI.

Boxplots are drawn with the box extending from the 25^th^ to 75^th^ percentiles, and the middle line plotted at the median. Whiskers reach the minimum and maximum values of the distribution. Details regarding the statistical tests employed, multiple hypothesis correction, and the use of repeated-measures statistical testing are outlined in the figure captions and Supplementary Table S1 and S2.

## Data and materials availability

Raw and processed data are available on the data platform DANDI: https://doi.org/10.48324/dandi.001635/0.251111.1144. All remaining data are available in the manuscript or the supplementary materials. For individual figures source data are provided with this paper.

## Code availability

The codes used for the processing and analysis of the raw data are made available as a GitHub repository - https://github.com/Wahl-lab/Panzeri_ContralesionalStroke

