## Supplementary Materials for "Contralesional activity reflects compensation, while brainstem detour pathways support skilled motor recovery after stroke"

for

### Supplementary Table S1, S2

### Supplementary Figures S1-S5

**Table S1**

| Panel | Statistical test | Post hoc test / multiple comparisons | Comparison | P-value |
|---|---|---|---|---|
| 1D | two-sample t-test | n.a. | Stroke vs. Stroke + training | 0,124 |
| 1E (middle) | Linear mixed effect model | Bonferroni | Early: sham vs. stroke | < 0.001 |
| | | | Early: sham vs. stroke + training | < 0.001 |
| | | | Late: sham vs. stroke | < 0.001 |
| | | | Late: sham vs. stroke + training | < 0.001 |
| 1E (right) | Linear mixed effect model | Bonferroni | Late: sham vs stroke + training | 0,001 |
| | | | Late: stroke vs. stroke + training | < 0.001 |
| 1F (left) | Linear mixed effect model | Bonferroni | Early: stroke vs. stroke + training | 0,025 |
| | | | Late: sham vs. stroke + training | 0,004 |
| 1F (middle) | Linear mixed effect model | Bonferroni | Early: sham vs. stroke | 0,001 |
| | | | Early: sham vs. stroke + training | 0,016 |
| | | | Late: sham vs. stroke | 0,001 |
| | | | Late: sham vs stroke + training | < 0.001 |
| 1F (right) | Linear mixed effect model | Bonferroni | Early: sham vs. stroke | < 0.001 |
| | | | Early: sham vs. stroke + training | 0,001 |
| | | | Late: sham vs. stroke | 0,042 |
| | | | Late: sham vs. stroke + training | < 0.001 |
| 1G | Spearman correlation | n.a. | | |
| 2D - E | Linear mixed effect models (one per ROI) | False discovery rate control over all comparisons. Benjamini-Hochberg correction | (MOp_R) Early: Sham vs. Stroke | < 0.001 |
| | | | (MOp_R) Early: Sham vs. Stroke + training | < 0.001 |
| | | | (MOp_R) Late: Sham vs. Stroke | 0,004 |
| | | | (MOp_R) Late: Sham vs. Stroke + training | 0,013 |
| | | | (MOp_L) Early: Sham vs. Stroke + training | 0,016 |
| | | | (MOp_L) Late: Stroke vs. Stroke + training | < 0.001 |
| | | | (MOs-lateral_R) Early: Sham vs. Stroke | < 0.001 |
| | | | (MOs-lateral_R) Early: Sham vs. Stroke + training | < 0.001 |
| | | | (MOs-lateral_L) Late: Stroke vs. Stroke + training | 0,048 |
| | | | (MOs-medial_R) Early: Sham vs. Stroke | 0,011 |
| | | | (MOs-medial_R) Early: Sham vs. Stroke + training | < 0.001 |
| | | | (MOs-medial_R) Late: Sham vs. Stroke | 0,020 |

| | | | | |
|---|---|---|---|---|
| | | | (MOs-medial_R) Late: Sham vs. Stroke + training | 0,026 |
| | | | (MOs-medial_L) Early: Sham vs. Stroke + training | 0,013 |
| | | | (RSP-anterior_R) Early: Sham vs. Stroke + training | 0,032 |
| | | | (RSP-anterior_L) Late: Stroke vs. Stroke + training | 0,027 |
| | | | (SSp-bfd_R) Early: Sham vs. Stroke | 0,011 |
| | | | (SSp-bfd_R) Late: Sham vs. Stroke | 0,012 |
| | | | (SSp-bfd_L) Late: Stroke vs. Stroke + training | 0,017 |
| | | | (SSp-ll_R) Early: Sham vs. Stroke | < 0.001 |
| | | | (SSp-ll_R) Early: Sham vs. Stroke + training | < 0.001 |
| | | | (SSp-ll_R) Late: Sham vs. Stroke | < 0.001 |
| | | | (SSp-ll_R) Late: Sham vs. Stroke + training | 0,015 |
| | | | (SSp-ll_L) Late: Stroke vs. Stroke + training | 0,004 |
| | | | (SSp-nose+mouth_R) Early: Sham vs. Stroke | 0,007 |
| | | | (SSp-nose+mouth_R) Early: Sham vs. Stroke + training | 0,002 |
| | | | (SSp-nose+mouth_L) Late: Sham vs. Stroke + training | 0,049 |
| | | | (SSp-nose+mouth_L) Late: Stroke vs. Stroke + training | 0,011 |
| | | | (SSp-tr_R) Early: Sham vs. Stroke | 0,018 |
| | | | (SSp-tr_R) Early: Sham vs. Stroke + training | 0,010 |
| | | | (SSp-tr_R) Late: Sham vs. Stroke | 0,017 |
| | | | (SSp-tr_L) Late: Stroke vs. Stroke + training | 0,016 |
| | | | (SSp-ul_R) Early: Sham vs. Stroke | < 0.001 |
| | | | (SSp-ul_R) Early: Sham vs. Stroke + training | < 0.001 |
| | | | (SSp-ul_R) Late: Sham vs. Stroke | < 0.001 |
| | | | (SSp-ul_R) Late: Sham vs. Stroke + training | < 0.001 |
| | | | (SSp-ul_L) Late: Sham vs. Stroke + training | 0,030 |
| | | | (SSp-ul_L) Late: Stroke vs. Stroke + training | < 0.001 |
| | | | (VIS-medial_R) Late: Stroke vs. Stroke + training | 0,004 |
| | | | (VIS-medial_L) Late: Stroke vs. Stroke + training | 0,016 |

| | | | | |
|---|---|---|---|---|
| | | | (VISa_R) Late: Sham vs. Stroke | 0,048 |
| | | | (VISa_R) Late: Stroke vs. Stroke + training | 0,037 |
| | | | (VISa_L) Late: Stroke vs. Stroke + training | 0,016 |
| | | | (VISp_R) Late: Stroke vs. Stroke + training | 0,037 |
| | | | (VISrl_L) Late: Stroke vs. Stroke + training | 0,040 |
| 2F | Spearman correlation | n.a. | | |
| 4A (top) | Linear mixed effect models (one per ROI) | False discovery rate control over all comparisons. Benjamini-Hochberg correction | (MOp_R) Early: Sham vs. Stroke | 0,002 |
| | | | (MOp_R) Early: Sham vs. Stroke + training | < 0.001 |
| | | | (MOp_R) Late: Sham vs. Stroke | 0,05 |
| | | | (MOp_R) Late: Sham vs. Stroke + training | 0,05 |
| | | | (MOs-lateral_R) Early: Sham vs. Stroke | 0,042 |
| | | | (MOs-medial_R) Early: Sham vs. Stroke | 0,038 |
| | | | (SSp-bfd_R) Early: Sham vs. Stroke | 0,017 |
| | | | (SSp-ll_R) Early: Sham vs. Stroke | < 0.001 |
| | | | (SSp-ll_R) Early: Sham vs. Stroke + training | < 0.001 |
| | | | (SSp-ll_R) Late: Sham vs. Stroke | 0,008 |
| | | | (SSp-ll_R) Late: Sham vs. Stroke + training | 0,009 |
| | | | (SSp-nosemouth_R) Early: Sham vs. Stroke + training | 0,037 |
| | | | (SSp-tr_R) Early: Sham vs. Stroke | 0,002 |
| | | | (SSp-tr_R) Early: Sham vs. Stroke + training | 0,007 |
| | | | (SSp-ul_R) Early: Sham vs. Stroke | < 0.001 |
| | | | (SSp-ul_R) Early: Sham vs. Stroke + training | < 0.001 |
| | | | (SSp-ul_R) Late: Sham vs. Stroke | 0,002 |
| | | | (SSp-ul_R) Late: Sham vs. Stroke + training | 0,002 |
| 4A (bottom) | Spearman correlation | n.a. | | |
| 4B (top) | Linear mixed effect models (one per ROI) | False discovery rate control over all comparisons. Benjamini-Hochberg correction | (MOp_R) Early: Sham vs. Stroke | 0,017 |
| | | | (MOp_R) Early: Sham vs. Stroke + training | 0,024 |
| | | | (MOp_L) Late: Stroke vs. Stroke + training | 0,013 |
| | | | (MOs-lateral_L) Late: Sham vs. Stroke + training | 0,041 |

| | | | | |
|---|---|---|---|---|
| | | | (MOs-lateral_L) Late: Stroke vs. Stroke + training | 0,013 |
| | | | (MOs-medial_R) Early: Sham vs. Stroke | 0,014 |
| | | | (MOs-medial_R) Early: Sham vs. Stroke + training | 0,028 |
| | | | (MOs-medial_R) Late: Sham vs. Stroke | 0,034 |
| | | | (MOs-medial_L) Late: Stroke vs. Stroke + training | 0,016 |
| | | | (RSP-anterior_L) Late: Stroke vs. Stroke + training | 0,034 |
| | | | (SSp-ll_R) Early: Sham vs. Stroke | 0,013 |
| | | | (SSp-ll_R) Early: Sham vs. Stroke + training | 0,016 |
| | | | (SSp-ll_L) Late: Stroke vs. Stroke + training | 0,016 |
| | | | (SSp-tr_R) Early: Sham vs. Stroke | 0,013 |
| | | | (SSp-tr_R) Early: Sham vs. Stroke + training | 0,023 |
| | | | (SSp-tr_R) Late: Stroke vs. Stroke + training | 0,041 |
| | | | (SSp-tr_L) Late: Stroke vs. Stroke + training | 0,016 |
| | | | (SSp-ul_R) Early: Sham vs. Stroke | 0,034 |
| | | | (SSp-ul_R) Early: Sham vs. Stroke + training | 0,041 |
| | | | (SSp-ul_L) Late: Stroke vs. Stroke + training | 0,047 |
| | | | (VISa_R) Late: Stroke vs. Stroke + training | 0,041 |
| | | | (VISa_L) Late: Stroke vs. Stroke + training | 0,034 |
| 4B (bottom) | Spearman correlation | n.a. | | |
| 4C (bottom) | Spearman correlation | n.a. | | |
| S1C-D | Spearman correlation | n.a. | | |
| S1F (right) | Linear mixed effect model | Bonferroni | Late: Sham vs. Stroke | 0,006 |
| | | | Late: sham vs. stroke + training | 0,047 |
| S2A-C | Spearman correlation | n.a. | | |
| S3B | Linear mixed effect models (one per ROI) | False discovery rate control over all comparisons. Benjamini-Hochberg correction | (MOp_R) Early: Sham vs. Stroke | < 0.001 |
| | | | (MOp_R) Early: Sham vs. Stroke + training | < 0.001 |
| | | | (MOp_R) Late: Sham vs. Stroke | 0,002 |
| | | | (MOp_R) Late: Sham vs. Stroke + training | 0,004 |

| | | | | |
|---|---|---|---|---|
| | | | (MOp_L) Early: Sham vs. Stroke | 0,047 |
| | | | (MOp_L) Early: Sham vs. Stroke + training | 0,013 |
| | | | (MOp_L) Late: Stroke vs. Stroke + training | 0,001 |
| | | | (MOs-lateral_R) Early: Sham vs. Stroke | < 0.001 |
| | | | (MOs-lateral_R) Early: Sham vs. Stroke + training | < 0.001 |
| | | | (MOs-lateral_R) Late: Sham vs. Stroke | 0,013 |
| | | | (MOs-lateral_L) Late: Stroke vs. Stroke + training | 0,025 |
| | | | (MOs-medial_R) Early: Sham vs. Stroke | 0,005 |
| | | | (MOs-medial_R) Early: Sham vs. Stroke + training | < 0.001 |
| | | | (MOs-medial_R) Late: Sham vs. Stroke | 0,005 |
| | | | (MOs-medial_R) Late: Sham vs. Stroke + training | 0,01 |
| | | | (MOs-medial_L) Early: Sham vs. Stroke | 0,047 |
| | | | (MOs-medial_L) Early: Sham vs. Stroke + training | 0,007 |
| | | | (RSP-anterior_R) Early: Sham vs. Stroke + training | 0,047 |
| | | | (SSp-bfd_R) Early: Sham vs. Stroke | 0,007 |
| | | | (SSp-bfd_R) Late: Sham vs. Stroke | 0,01 |
| | | | (SSp-ll_R) Early: Sham vs. Stroke | 0,002 |
| | | | (SSp-ll_R) Early: Sham vs. Stroke + training | < 0.001 |
| | | | (SSp-ll_R) Late: Sham vs. Stroke | 0,001 |
| | | | (SSp-ll_R) Late: Sham vs. Stroke + training | 0,009 |
| | | | (SSp-ll_L) Late: Stroke vs. Stroke + training | 0,024 |
| | | | (SSp-nosemouth_R) Early: Sham vs. Stroke | 0,003 |
| | | | (SSp-nosemouth_R) Early: Sham vs. Stroke + training | < 0.001 |
| | | | (SSp-nosemouth_L) Late: Stroke vs. Stroke + training | 0,001 |
| | | | (SSp-tr_R) Early: Sham vs. Stroke | 0,024 |
| | | | (SSp-tr_R) Early: Sham vs. Stroke + training | 0,018 |
| | | | (SSp-tr_R) Late: Sham vs. Stroke | 0,013 |
| | | | (SSp-tr_L) Late: Stroke vs. Stroke + training | 0,047 |

| | | | | |
|---|---|---|---|---|
| | | | (SSp-ul_R) Early: Sham vs. Stroke | < 0.001 |
| | | | (SSp-ul_R) Early: Sham vs. Stroke + training | < 0.001 |
| | | | (SSp-ul_R) Late: Sham vs. Stroke | < 0.001 |
| | | | (SSp-ul_R) Late: Sham vs. Stroke + training | < 0.001 |
| | | | (SSp-ul_L) Late: Stroke vs. Stroke + training | 0,004 |
| | | | (VISa_R) Late: Sham vs. Stroke | 0,029 |
| | | | (VISa_L) Late: Stroke vs. Stroke + training | 0,047 |
| | | | (VISrl_R) Late: Sham vs. Stroke | 0,047 |
| S3D | Linear mixed effect models (one per ROI) | False discovery rate control over all comparisons. Benjamini-Hochberg correction | (MOp_R) Early: Sham vs. Stroke | 0,002 |
| | | | (MOp_R) Early: Sham vs. Stroke + training | < 0.001 |
| | | | (MOp_R) Late: Sham vs. Stroke | 0,011 |
| | | | (MOp_R) Late: Sham vs. Stroke + training | 0,022 |
| | | | (MOp_L) Late: Stroke vs. Stroke + training | 0,005 |
| | | | (MOs-lateral_R) Early: Sham vs. Stroke | < 0.001 |
| | | | (MOs-lateral_R) Early: Sham vs. Stroke + training | < 0.001 |
| | | | (MOs-medial_R) Early: Sham vs. Stroke | 0,002 |
| | | | (MOs-medial_R) Early: Sham vs. Stroke + training | < 0.001 |
| | | | (MOs-medial_R) Late: Sham vs. Stroke | 0,014 |
| | | | (MOs-medial_R) Late: Sham vs. Stroke + training | 0,027 |
| | | | (MOs-medial_L) Early: Sham vs. Stroke + training | 0,014 |
| | | | (SSp-bfd_R) Late: Sham vs. Stroke | 0,046 |
| | | | (SSp-ll_R) Early: Sham vs. Stroke | 0,005 |
| | | | (SSp-ll_R) Early: Sham vs. Stroke + training | < 0.001 |
| | | | (SSp-ll_R) Late: Sham vs. Stroke | 0,004 |
| | | | (SSp-ll_R) Late: Sham vs. Stroke + training | 0,037 |
| | | | (SSp-ll_L) Late: Stroke vs. Stroke + training | 0,024 |
| | | | (SSp-nosemouth_R) Early: Sham vs. Stroke | 0,037 |
| | | | (SSp-nosemouth_R) Early: Sham vs. Stroke + training | 0,009 |

| | | | | |
|---|---|---|---|---|
| | | | (SSp-nosemouth_L) Late: Stroke vs. Stroke + training | 0,024 |
| | | | (SSp-tr_R) Early: Sham vs. Stroke + training | 0,037 |
| | | | (SSp-tr_R) Late: Sham vs. Stroke | 0,041 |
| | | | (SSp-tr_L) Late: Stroke vs. Stroke + training | 0,024 |
| | | | (SSp-ul_R) Early: Sham vs. Stroke | < 0.001 |
| | | | (SSp-ul_R) Early: Sham vs. Stroke + training | < 0.001 |
| | | | (SSp-ul_R) Late: Sham vs. Stroke | < 0.001 |
| | | | (SSp-ul_R) Late: Sham vs. Stroke + training | 0,001 |
| | | | (SSp-ul_L) Late: Stroke vs. Stroke + training | 0,014 |
| | | | (VIS-medial_R) Late: Stroke vs. Stroke + training | 0,014 |
| | | | (VIS-medial_L) Late: Stroke vs. Stroke + training | 0,016 |
| | | | (VISa_L) Late: Stroke vs. Stroke + training | 0,022 |
| | | | (VISp_L) Late: Stroke vs. Stroke + training | 0,037 |
| | | | (VISrl_L) Late: Stroke vs. Stroke + training | 0,038 |
| S4A | Linear mixed effect models (one per ROI) | False discovery rate control over all comparisons. Benjamini-Hochberg correction | (MOp_R) Early: Sham vs. Stroke | < 0.001 |
| | | | (MOp_R) Early: Sham vs. Stroke + training | < 0.001 |
| | | | (MOp_R) Late: Sham vs. Stroke | 0,004 |
| | | | (MOp_R) Late: Sham vs. Stroke + training | 0,01 |
| | | | (MOp_L) Early: Sham vs. Stroke + training | 0,04 |
| | | | (MOp_L) Late: Stroke vs. Stroke + training | < 0.001 |
| | | | (MOs-lateral_R) Early: Sham vs. Stroke | < 0.001 |
| | | | (MOs-lateral_R) Early: Sham vs. Stroke + training | < 0.001 |
| | | | (MOs-lateral_L) Late: Stroke vs. Stroke + training | 0,04 |
| | | | (MOs-medial_R) Early: Sham vs. Stroke | 0,003 |
| | | | (MOs-medial_R) Early: Sham vs. Stroke + training | < 0.001 |
| | | | (MOs-medial_R) Late: Sham vs. Stroke | 0,005 |
| | | | (MOs-medial_R) Late: Sham vs. Stroke + training | 0,004 |

| | | | | |
|---|---|---|---|---|
| | | | (MOs-medial_L) Early: Sham vs. Stroke + training | 0,013 |
| | | | (MOs-medial_L) Late: Stroke vs. Stroke + training | 0,043 |
| | | | (RSP-anterior_R) Early: Sham vs. Stroke | 0,036 |
| | | | (RSP-anterior_R) Early: Sham vs. Stroke + training | 0,013 |
| | | | (RSP-anterior_L) Late: Stroke vs. Stroke + training | 0,027 |
| | | | (RSP-posterior_R) Late: Stroke vs. Stroke + training | 0,013 |
| | | | (RSP-posterior_L) Late: Stroke vs. Stroke + training | 0,025 |
| | | | (SSp-bfd_R) Early: Sham vs. Stroke | 0,005 |
| | | | (SSp-bfd_R) Late: Sham vs. Stroke | 0,003 |
| | | | (SSp-bfd_L) Late: Stroke vs. Stroke + training | 0,011 |
| | | | (SSp-ll_R) Early: Sham vs. Stroke | < 0.001 |
| | | | (SSp-ll_R) Early: Sham vs. Stroke + training | < 0.001 |
| | | | (SSp-ll_R) Late: Sham vs. Stroke | < 0.001 |
| | | | (SSp-ll_R) Late: Sham vs. Stroke + training | 0,004 |
| | | | (SSp-ll_L) Late: Stroke vs. Stroke + training | 0,001 |
| | | | (SSp-nosemouth_R) Early: Sham vs. Stroke | 0,005 |
| | | | (SSp-nosemouth_R) Early: Sham vs. Stroke + training | 0,003 |
| | | | (SSp-nosemouth_L) Late: Stroke vs. Stroke + training | 0,01 |
| | | | (SSp-tr_R) Early: Sham vs. Stroke | 0,001 |
| | | | (SSp-tr_R) Early: Sham vs. Stroke + training | 0,001 |
| | | | (SSp-tr_R) Late: Sham vs. Stroke | 0,002 |
| | | | (SSp-tr_R) Late: Stroke vs. Stroke + training | 0,045 |
| | | | (SSp-tr_L) Late: Stroke vs. Stroke + training | 0,004 |
| | | | (SSp-ul_R) Early: Sham vs. Stroke | < 0.001 |
| | | | (SSp-ul_R) Early: Sham vs. Stroke + training | < 0.001 |
| | | | (SSp-ul_R) Late: Sham vs. Stroke | < 0.001 |
| | | | (SSp-ul_R) Late: Sham vs. Stroke + training | < 0.001 |
| | | | (SSp-ul_L) Late: Sham vs. Stroke + training | 0,042 |

| | | | | |
|---|---|---|---|---|
| | | | (SSp-ul_L) Late: Stroke vs. Stroke + training | < 0.001 |
| | | | (VIS-medial_R) Late: Sham vs. Stroke | 0,036 |
| | | | (VIS-medial_R) Late: Stroke vs. Stroke + training | < 0.001 |
| | | | (VIS-medial_L) Late: Stroke vs. Stroke + training | 0,003 |
| | | | (VISa_R) Early: Sham vs. Stroke | 0,015 |
| | | | (VISa_R) Late: Sham vs. Stroke | 0,005 |
| | | | (VISa_R) Late: Stroke vs. Stroke + training | 0,008 |
| | | | (VISa_L) Late: Stroke vs. Stroke + training | 0,005 |
| | | | (VISp_R) Late: Stroke vs. Stroke + training | 0,03 |
| | | | (VISp_L) Late: Stroke vs. Stroke + training | 0,01 |
| | | | (VISrl_R) Late: Sham vs. Stroke | 0,036 |
| | | | (VISrl_R) Late: Stroke vs. Stroke + training | 0,027 |
| | | | (VISrl_L) Late: Stroke vs. Stroke + training | 0,005 |

**Supplementary Table 1.** Statistical tests, post-hoc tests (if applicable) and exact p-values (3 significant figures) of significant comparisons used in figure panels. Region of interest (ROI) names are suffixed with "_L" for the left hemisphere shown in the panels, and "_R" for the right hemisphere shown in the panels. ROI abbreviations are fully named in supplementary table 2.

**Table S2**

| **Area abbreviation** | **Area description** |
|---|---|
| MOp | Primary motor cortex (M1) |
| MOs-lateral | Anterior-lateral portion of the secondary motor cortex (M2) |
| MOs-medial | Posterior-medial portion of the secondary motor cortex (M2) |
| SSp-ll | Somatosensory cortex - hindlimb |
| SSp-ul | Somatosensory cortex - forelimb |
| SSp-nosemouth | Merged ROI from mouth and nose somatosensory ROIs |
| SSp-bfd | Somatosensory cortex - barrel cortex |
| SSp-tr | Somatosensory cortex - trunk |
| RSp-anterior | Anterior part of the retrosplenial cortex |
| RSP-posterior | Posterior part of the retrosplenial cortex |
| VISp | Primary visual cortex (V1) |
| VIS-medial | Medial visual areas |
| VISa | Posterior parietal cortex - area "a" ("PPC-a") |
| VISrl | Posterior parietal cortex - area "rl" ("PPC-rl") |

**Supplementary Table 2**. List of regions of interest names and abbreviations, based on the Allen Brain Atlas.

**Supplementary Figure 1**

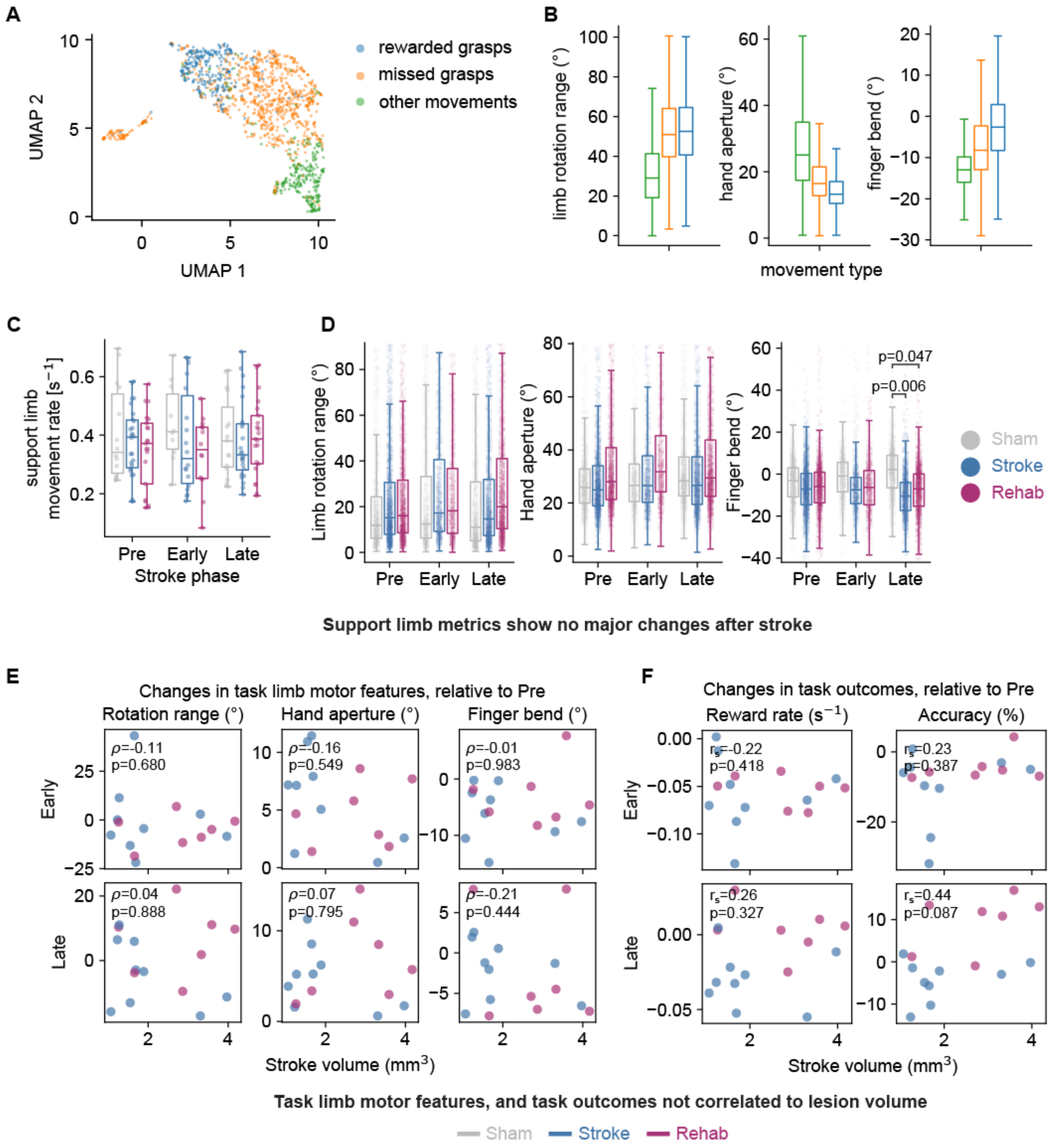


**Supplementary Figure 1. Further fine motor skill quantifications. A.** UMAP embedding of all pre-stroke grasps from one mouse, color coded by the type of movement (n=1497 grasps, n=1 mice). **B.** Comparison of fine motor features across different movement types. Data from all detected pre-stroke movements (n=28033 grasps, n=21 mice). **C.** Comparison of the rate of support limb movements (number of movements per second) between different cohorts. Datapoints are individual sessions (n=163 sessions, n=21 mice). **D.** Comparison of support limb fine motor features across cohorts. Datapoints represent average support limb fine motor feature during a rewarded grasp (n=14115 rewarded grasps, n=21 mice).**E.** Spearman correlations between task limb motor features and stroke volume. Individual datapoints represent average motor features for each mouse in the given experimental phase, relative to pre-stroke (n=16 mice). **F.** Spearman correlations between task-performance metrics (reward rate and accuracy), and lesion size. Datapoints represent averages over the experimental phase for each mouse (n=16 mice). In **C.** and **D.** statistical comparisons are performed using linear mixed effect models (one model per plot) and p-values are adjusted per-model for multiple comparisons (post-hoc Bonferroni correction).

**Supplementary Figure 2**

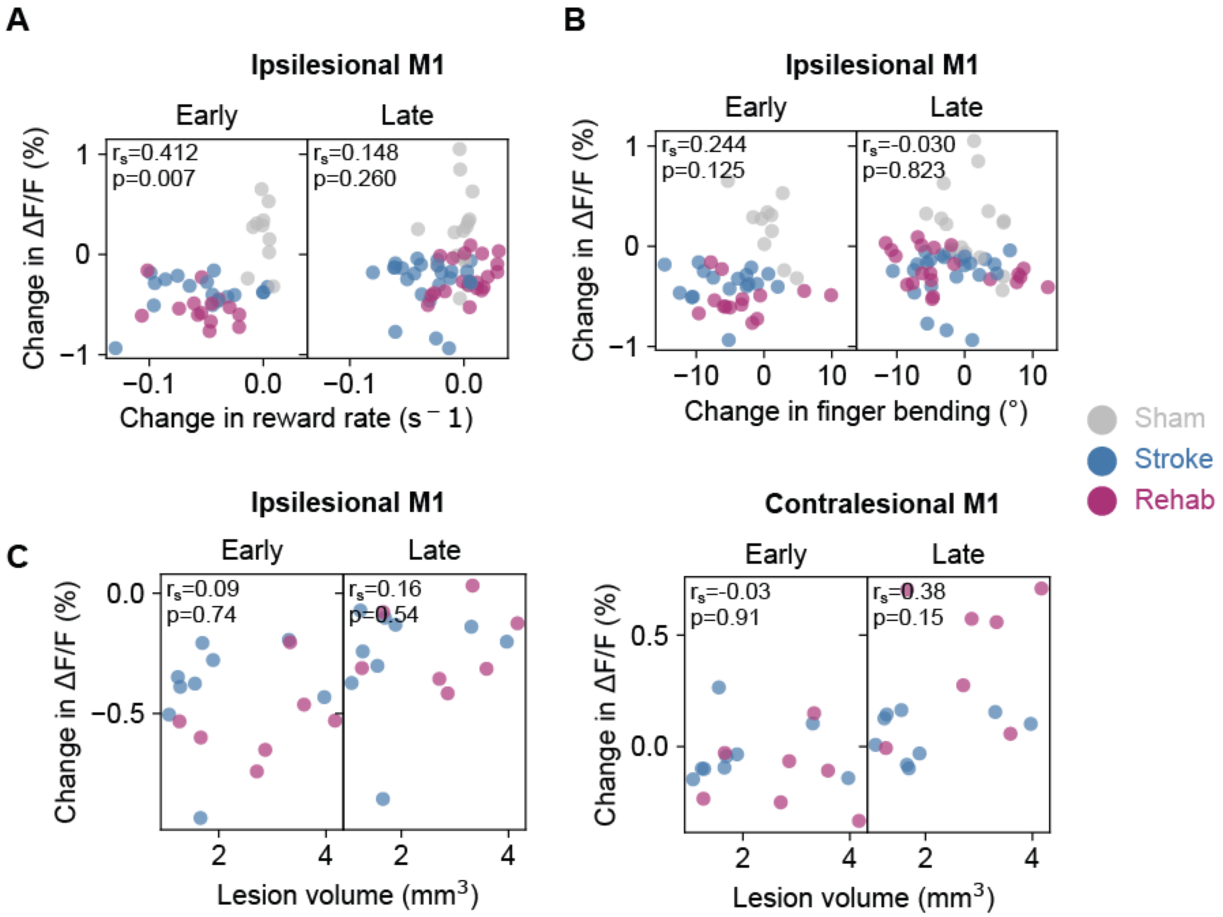


**Supplementary Figure 2. Correlations of activity with behavioral parameters and lesion sizes. A.** Spearman correlation between change in $\Delta F/F_0$ of the ipsilesional M1 and session reward rate. Individual datapoints sessions averages over rewarded grasps (n=163 sessions, n=21 mice) and are relative to pre-stroke baseline. **B.** Same as for **A.** but correlating the activity with the session-average finger bending fine motor parameters. **C.** Spearman correlations between change in cortical activity (averaged for all grasps within session) and lesion volume. Left: ipsilesional M1, right: contralesional M1. Individual datapoints average over the given experimental phase (n=16 mice) and are relative to pre stroke baseline.

### Supplementary Figure 3

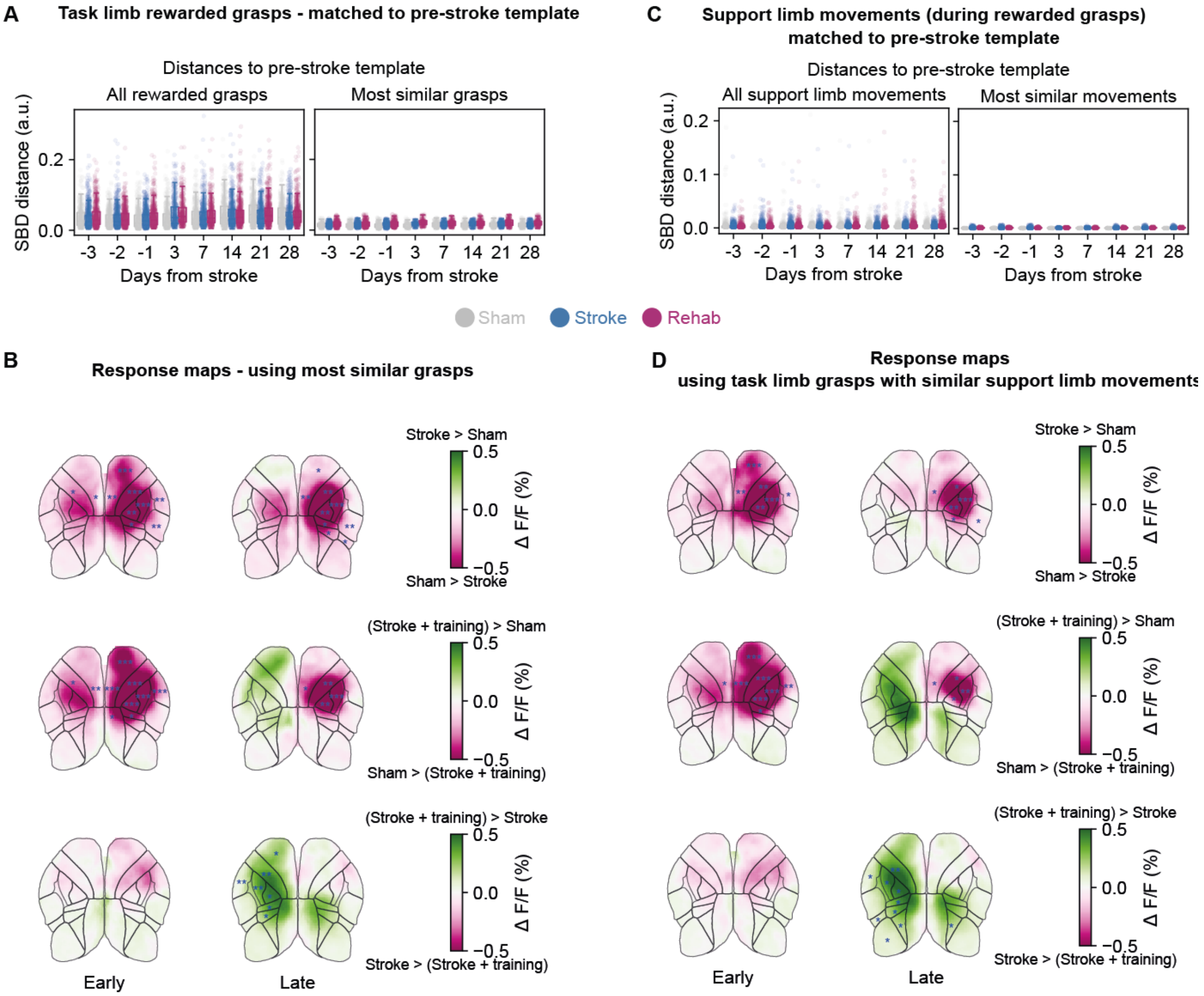


**Supplementary Figure 3. Increase in contralesional activity is not driven by changes in task or support limb trajectories. A.** Shape-based distances (SBD) between individual task-limb grasps their pre-stroke template grasp (calculated per mouse). Datapoints represent individual grasps. Left: all rewarded grasps (n=14115 grasps, n=21 mice). Right: grasps that are classified as similar to pre-stroke template (n=6073 grasps, n=21 mice). **B.** Differences in $\Delta F/F_0$ response maps between groups, using the subset of similar grasps. Maps obtained by averaging response-window activity over grasps (n=6073 grasps, n=21 mice). Data is normalized to pre-stroke baseline. Blue overlays represent results of statistical testing. **C.** Same as **A.** but matching support-limb trajectories (during rewarded task-limb grasps). Left: all support limb movements (n=14115 grasps, n=21 mice), right: subset of support limb movements classified as similar (n=6411 grasps, n=21 mice). **D.** same as **B.** but using the subset of task-limb that exhibit similar support-limb movements. In **B.** and **D.** statistical comparisons are computed using linear mixed-effect models (one per ROI) and p-values are adjusted by controlling the false discovery rate (Benjamini-Hochberg correction). Blue asterisks indicate significances for the corresponding outlined ROI: *p<0.05, **p<0.01, *** p<0.001.

**Supplementary Figure 4**

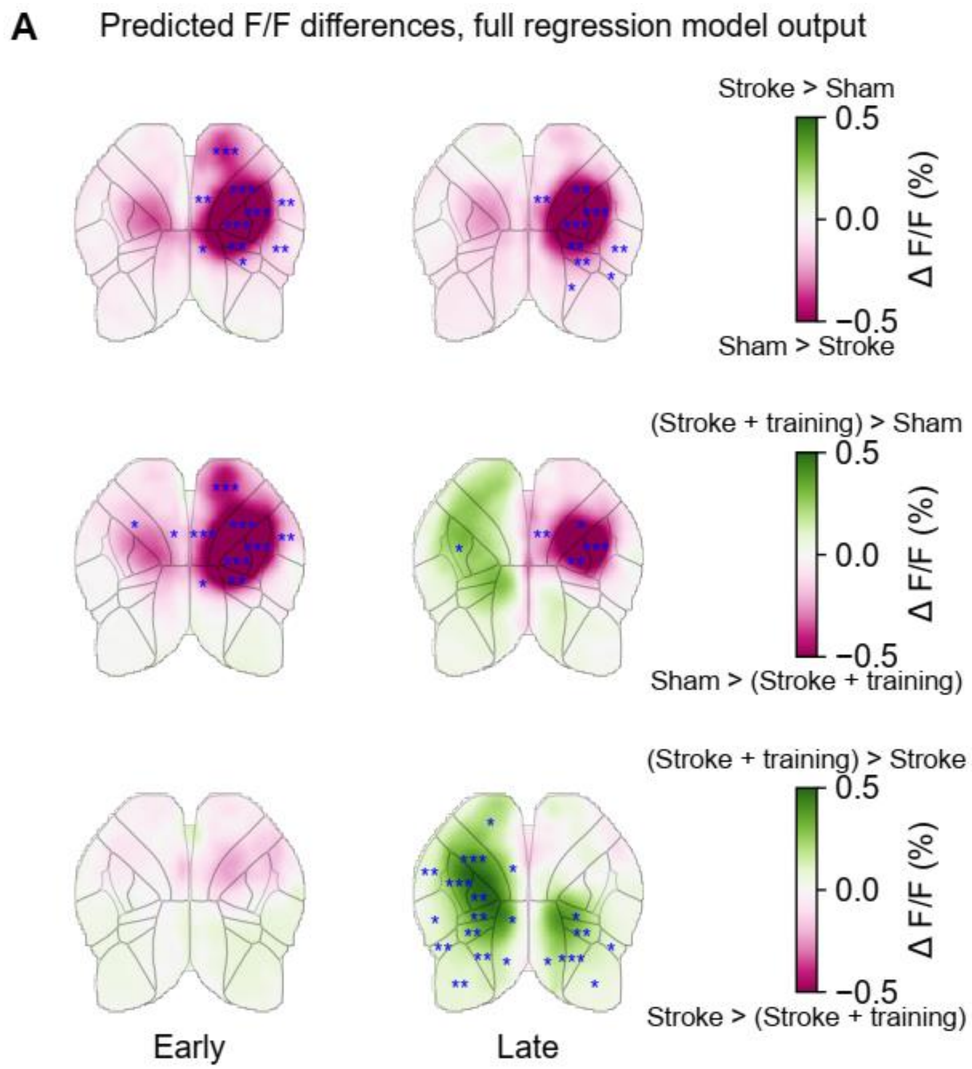


**Supplementary Figure 4. Outputs of the full ridge regression models capture the increase in contralesional activity seen in the widefield imaging data. A.** Differences in $\Delta F/F_0$ predicted by the full ridge regression model, between cohorts at different post-stroke phases. Datapoints are response window averaged $\Delta F/F_0$, over individual grasps (n=102 sessions, n=21 mice). Statistical comparisons are computed using linear mixed-effect models (one per ROI) and p-values are adjusted for multiple comparisons by controlling the false discovery rate (Benjamini-Hochberg correction). Blue asterisks indicate significances for the corresponding outlined ROI: *p<0.05, **p<0.01, *** p<0.001.

**Supplementary Figure 5**

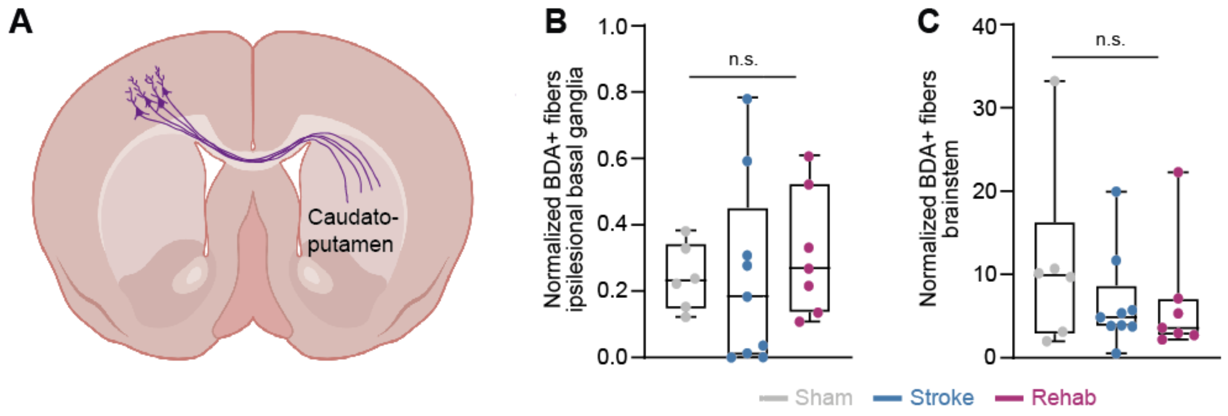


**Supplementary Figure 5. A.** Scheme revealing BDA+ fibers projecting from the contralesional hemisphere to the basal ganglia. **B.** Normalized BDA+ fiber count in the ipsilesional basal ganglia were compared for the different experimental groups (Sham n=6, Stroke n=9, Stroke + training group n=7). **C.** Total BDA+ fiber density in the brain in the three experimental groups. For **B.** and **C.** statistical comparison was performed with a Kruskal-Wallis test setting a significance level of $p<0.05$.
